# A fine kinetic balance of interactions directs transcription factor hubs to genes

**DOI:** 10.1101/2024.04.16.589811

**Authors:** Samantha Fallacaro, Apratim Mukherjee, Puttachai Ratchasanmuang, Joseph Zinski, Yara I Haloush, Kareena Shankta, Mustafa Mir

## Abstract

Eukaryotic gene regulation relies on the binding of sequence-specific transcription factors (TFs). TFs bind chromatin transiently yet occupy their target sites by forming high-local concentration microenvironments (hubs and condensates) that increase the frequency of binding. Despite their ubiquity, such microenvironments are difficult to study in endogenous contexts due to technical limitations. Here, we use live embryo light-sheet imaging, single-molecule tracking, and genomics to overcome these limitations and investigate how hubs are localized to target genes to drive TF occupancy and transcription. By examining mutants of a hub-forming TF, Zelda, in *Drosophila* embryos, we find that hub formation propensity, spatial distributions, and temporal stabilities are differentially regulated by DNA binding and disordered protein domains. We show that hub localization to genomic targets is driven by a finely-tuned kinetic balance of interactions between proteins and chromatin, and hubs can be redirected to new genomic sites when this balance is perturbed.

## INTRODUCTION

Eukaryotic gene regulation is orchestrated by sequence-specific transcription factors (TFs) that selectively bind DNA sequence motifs and regulate transcriptional activity in concert with a multitude of binding partners and cofactors. The ability to measure TF-chromatin interaction kinetics in vivo using single-molecule tracking has revealed that most TFs exhibit surprisingly short residence times on chromatin, on the order of just tens of seconds or less^1–4^. These short residence times raised a kinetic conundrum regarding how transcription factors can robustly occupy their sites to drive minutes-long bursts of transcriptional activity, especially at low nuclear TF concentrations, where binding frequencies to any given site are expected to be lower than at higher nuclear concentrations. A resolution to this conundrum has come from a series of observations suggesting that instead of TF target occupancy being regulated by the stability of DNA binding, target occupancy may be dominated by increasing the frequency of binding events^5–17^. This increase in frequency is achieved by accumulating TFs around their targets in high local-concentration microenvironments referred to as condensates, implying liquid-like properties and phase separation based formation, or hubs, when agnostic to the mechanisms of formation^5,8,9,11,18–22^.

The formation of transcriptional hubs and condensates is often driven by selective but low-affinity interactions between the intrinsically disordered regions (IDRs) of TFs, outside of their sequence-specific and structured DNA binding domains (DBDs)^21,23–25^. While TFs are canonically described as modular, with the DBD dictating target selectivity and the disordered activation or interaction domains imparting function^26–28^, recent work suggests that DBDs and IDRs synergistically confer target site selection and occupation^6,29,30^. The mechanisms by which IDRs and DBDs work together to localize hubs, and the TFs within them, to their target sites are unknown. Our limited understanding may stem from the fact that most studies on condensate formation are performed in exogenous or overexpressed contexts or are focused on more stable assemblies that form at particular loci.

We and others have previously described transient multifactor hubs formed by the pioneer transcription factor Zelda in *Drosophila* embryos^18,31^. Zelda facilitates the binding of a majority of developmental TFs to activate thousands of genes during zygotic genome activation^32–34^. Incorporation into Zelda hubs allows morphogenic TFs to robustly occupy target sites even at low nuclear concentrations^5,18,35^. Here, we examine the underlying determinants of how Zelda hubs form and localize to targets through single-molecule tracking, lattice light-sheet imaging, genomics, and mutagenesis experiments.

## RESULTS

### Zelda hub formation results from a combination of DNA binding and IDR mediated interactions

Zelda (ZLD) is maternally deposited in Drosophila embryos as a 1596 amino acid protein with a cluster of four C_2_H_2_ zinc-fingers near its C-terminus (ZFs 3-6) and two upstream zinc-fingers (ZFs 1-2), with much of the remainder of it predicted to be intrinsically disordered (Fig. 1A)^36^. The C-terminal zinc-finger cluster (ZFs 3-6) comprises Zelda’s DNA binding domain (DBD) which recognizes its cognate binding motifs in the early embryo^37^. Mutations to any of ZFs 4-6 ablates specific binding to cognate motifs in vitro, prevents transcriptional activation in luciferase assays, and leads to developmental arrest during gastrulation^37^. The upstream zinc-fingers (ZFs 1-2) are not critical for Zelda’s function or site-specific targeting in the blastoderm, homozygous ZF1 mutants survive to adulthood while ZF2 mutants gastrulate but arrest later in embryogenesis^36^. We first sought to determine the roles of DNA binding and IDR mediated interactions in Zelda hub formation. We generated embryos expressing wildtype or mutant forms of Zelda fused to mNeonGreen for light-sheet imaging. We imaged the following mutants (Movie 1): (1) ZF1-6/+: All six ZFs scrambled (heterozygous only), (2) ΔIDR/+: Retaining only its DBD (heterozygous only; homozygous embryos die before visible nuclei), and (3) ZF5: A mutation to the two zinc-binding cysteines in the fifth zinc finger to serines which abolishes Zelda’s ability to recognize its canonical motifs in electromobility shift assays^37^ (Figs. 1B, S1).

**Fig 1.**
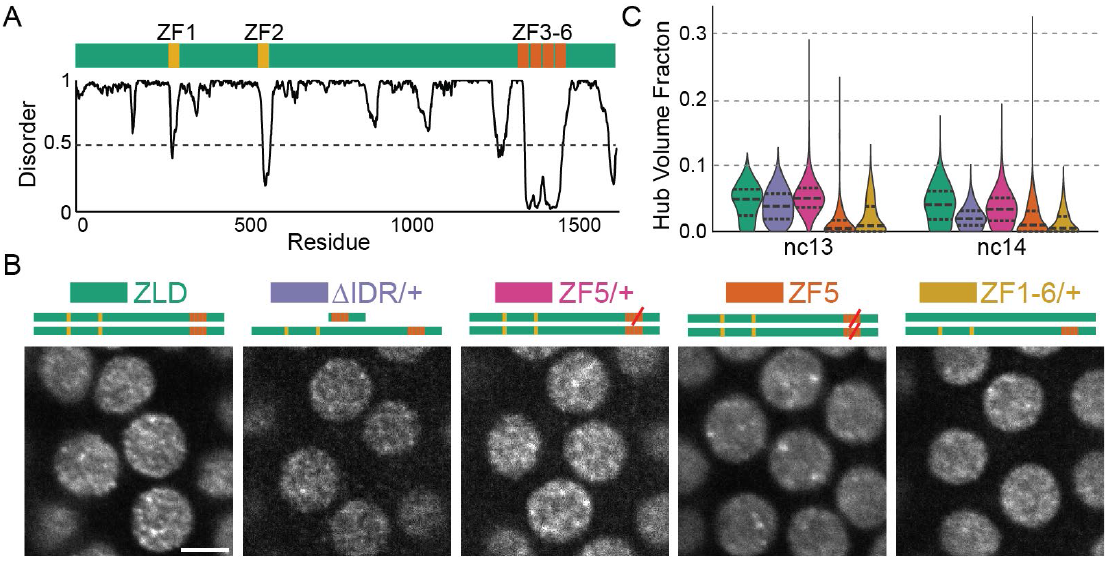
Zelda hub formation is differentially tuned by IDR and DNA binding interactions binding. **(A)** Domain map of Zelda indicating its DNA Binding Domain (ZF3-6). Graph shows predicted disorder score (from Metapredict^51^). **(B)** Light-sheet images (contrast adjusted) of mNeonGreen-ZLD, ZF5, ΔIDR/+, or ZF1-6/+ in the middle of interphase during nuclear cycle (nc) 13. Scale bar is 5 µm. **(C)** Fraction of nuclear volume occupied by hubs in each condition.

We find that when expressed heterozygously, ΔIDR molecules form hubs similar in appearance and occupy comparable a similar fraction of the nuclear volume as wildtype ZLD, despite a lower expression level (Figs. 1C, S2). ZF1-6 molecules incorporate less efficiently into hubs but their patterning is also similar in appearance to wildtype ZLD. Heterozygous ZF5 embryos also display similar hubs as wildtype ZLD but incorporate more efficiently into hubs than ZF1-6. Together these observations indicate that in the presence of wild type ZLD the DBD and IDR domains alone are both sufficient, but neither are necessary for hub incorporation, and that DNA binding ability modulates hub incorporation propensity. In stark contrast, when expressed homozygously, ZF5 forms hubs that appear larger and more static and the granular hub pattern present in the heterozygous conditions is lost (Figs. 1B-C, S2, and Movie 1). We speculated that homozygous ZF5 hubs may be forming at non-target genes driven by protein-protein interactions and thus serve as a model to provide insights on how hub targeting to genomic locations occurs independently of DBD-driven canonical motif recognition.

### A Zelda DBD mutant (ZF5) relocalizes to new genomic targets

To assess the effects of the ZF5 mutation on Zelda’s genomic localization and function we performed RNA-seq and CUT&RUN in wildtype and ZF5 homozygous and heterozygous embryos (Figs. 2, S3, Tables S1-S2). We found that ZF5 exhibits a completely divergent pattern of binding than ZLD when expressed homozygously (Fig. 2A). Consistent with Zelda’s role as an activator, we found a large number of down-regulated genes in ZF5 embryos (Figs. 2B, Table S3)^34,38^. However, we were intrigued to also find a set of significantly up-regulated genes in ZF5 embryos of which only 8.9% are also upregulated in Zelda null embryos (Fig. S3D-E)^34^. This differential activity suggests that the genes up-regulated in ZF5 embryos might be due to a relocalization of ZF5 to these genes. Consistent with this hypothesis CUT&RUN data shows that ZF5 localizes to the Transcription Start Sites (TSSs) and gene bodies of up-regulated genes (2.1-fold increase), and conversely, ZF5 is not binding at the TSSs and gene bodies of genes that are down-regulated in ZF5 embryos (9.2-fold decrease) (Fig. 2C-D). When expressed heterozygously, ZF5 doesn’t localize to these up-regulated genes and in fact exhibits diminished overall genomic association (Fig. 2B). ATAC-seq analysis shows that homozygous expressed ZF5’s relocalization is also correlated with a 2-fold increase in accessibility at up-regulated genes. This gain in accessibility at up-regulated genes is also present in ΔZLD embryos (Figs. 2E-F, S4 and Table S3). Together these data suggest that ZF5 relocalization is driven by the activation of aberrant transcriptional programs from the loss of ZLD binding to its cognate motifs, which leads to an increase in accessibility at ectopic sites. Contrary to recent observations on regions outside of a TFs DBD driving site-specific occupation at canonical target sites^29,39^, we find that the ZF5 mutation causes Zelda to broadly relocalize to new sites suggesting that a more complex interplay between factors drives the genomic localization of ZF5.

**Fig 2.**
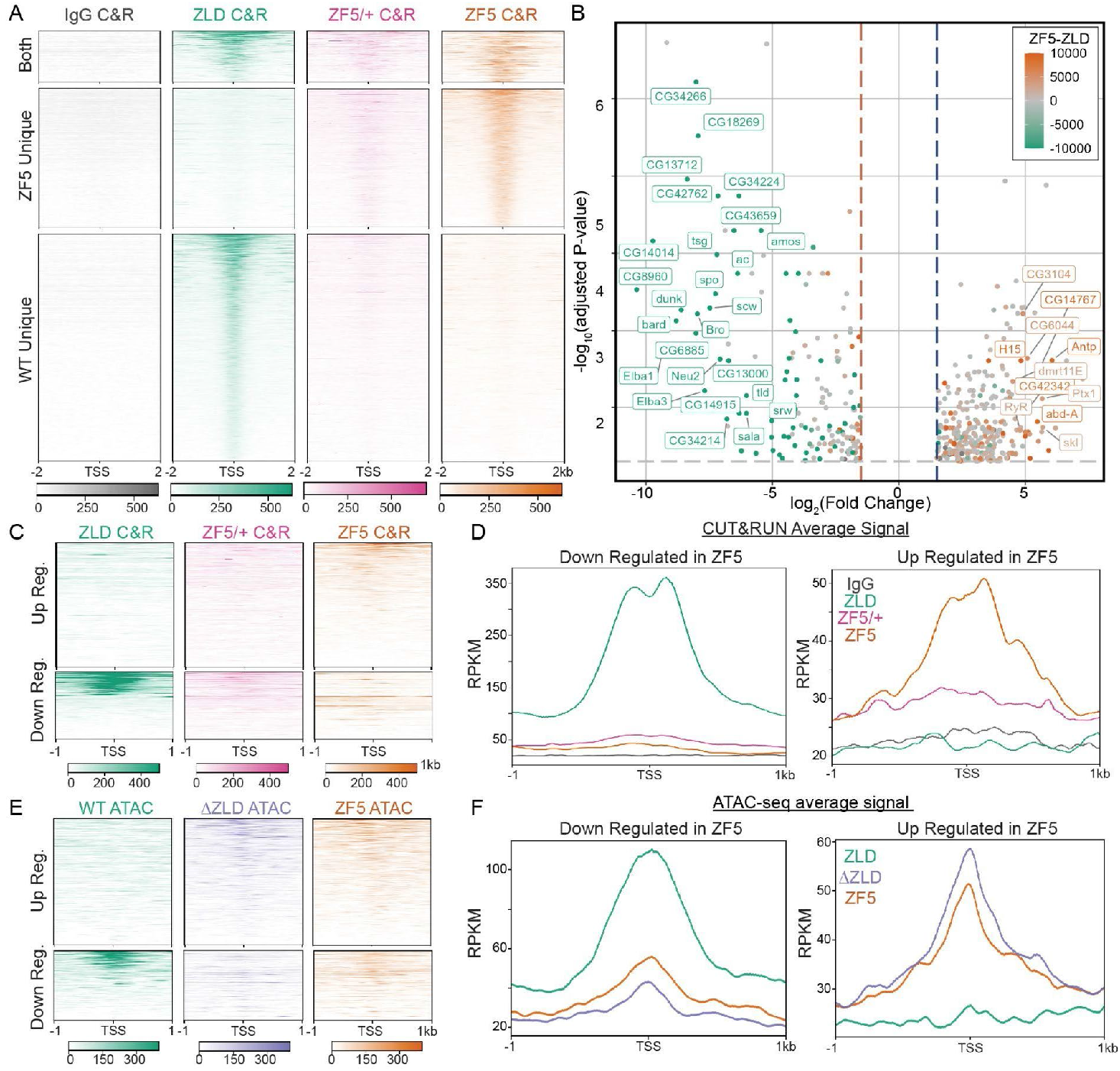
Zelda DNA binding domain mutation causes a redistribution to new genomic targets. **(A)** Heatmap of ZLD, ZF5 heterozygotes (ZF5/+), and ZF5 homozygous CUT&RUN peaks at TSSs separated by whether they are present in WT only (2262 genes), ZF5 only (1478 genes), or both (653 genes). **(B)** Volcano plot of genes up- and down-regulated in ZF5 vs ZLD via RNA-seq. Points are colored by the difference in CUT&RUN (C&R) reads between ZLD and ZF5 within 2kb of the TSS. Genes with an adjusted P-value <0.01, Log_2_ Fold Change > 4, and an absolute difference in RPKM > 5000 are labeled. The gray dotted line marks adj. P-value = 0.05, blue and red dotted lines are ±1.5 Fold Change. **(C-D)** Heatmaps and average profiles of CUT&RUN reads around the TSSs of differentially up and down regulated genes. **(E-F)** Heatmaps and average profiles of ATAC-seq reads around the TSSs of differentially up- and down-regulated genes.

### Perturbation to Zelda’s DBD alters its chromatin interaction and target search strategy

To determine how the ZF5 mutation influences its interaction kinetics with chromatin, we used long-exposure time (500 msec/frame) single-molecule tracking^5,18^ and found that the residence times of ZF5 molecules are reduced to just ~1 second compared to ~5 seconds for ZLD (Movie 2, Figs. 3A-B, and S5A). Together, the genomics and residence time data show that despite its impaired ability to stably interact with DNA, ZF5 can still occupy genes and drive expression.

**Fig 3.**
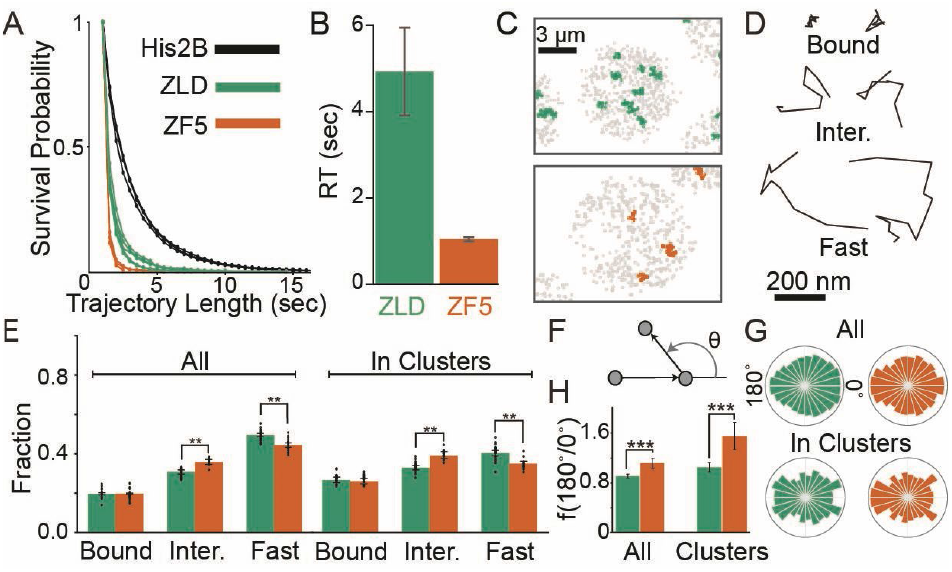
Mutation to Zelda’s DNA Binding Domain alters its chromatin interaction kinetics and target search strategy. **(A)** Raw survival probabilities of single-molecule trajectories at 500 msec/frame from at least three individual embryos for mEos3.2-ZLD, mEos3.2-ZF5, and H2B-mEos3.2. **(B**) Residence times after correction for photobleaching for mEos3.2-ZLD (4.93±1.01 sec) and mEos3.2-ZF5 (1.04±0.05 sec). **(C)** Representative maps of average trajectory positions (in gray) in nuclear cycle 14 from tracks accumulated over 80 sec at 10 msec/frame for ZLD (green) and ZF5 (orange). **(D)** Example trajectories from each kinetic category. **(E)** Bound, intermediate, and free fractions overall (left) and inside clusters (right). Error bars represent standard deviations from resampling analysis (see additional statistical analyses in Table S4). Points show data from 23 and 14 individual ZLD and ZF5 movies respectively, from at least three embryos each (n=70,827 tracks overall and 14,156 in clusters for ZLD and n=40,307 overall and 3,365 in clusters for ZF5, see Table S4 for data). **(F)** Angles are calculated between three consecutive displacements in non-bound tracks. **(G)** Distribution of angles overall (top) and in clusters (bottom). **(H)** Fold anisotropy overall (left) and in clusters (right). n=9,806 overall and 1,673 in clusters for ZLD and 4,609 overall and 995 in clusters for ZF5. Error bars represent standard deviations from resampling analysis.

We next used fast single-molecule tracking (10 msec/frame) (Movies 3-4) to investigate how the ZF5 mutation affects Zelda’s nuclear exploration kinetics. We found that single-molecule trajectories for both ZLD and ZF5 are spatially clustered, though consistent with our volumetric imaging in Fig. 1, the number of clusters is reduced for ZF5 (Figs. 3C and S5B). ZLD and ZF5 molecules exhibit a range of kinetic behaviors inside and out of clusters. To quantify these kinetics we inferred the diffusion coefficients of trajectories using state-arrays^40^ and assigned tracks into kinetic bins of chromatin bound, intermediate, and fast moving molecules (Figs. 3D and S5C-F). Surprisingly, we found that despite the loss in residence time, the fraction of bound molecules does not change in ZF5 when compared to ZLD. (Fig. 3E, Table S4). In contrast, the intermediate fraction is significantly higher in ZF5 than ZLD throughout the nucleus and this difference is more pronounced inside clusters.

The increase in the ZF5 intermediate kinetic population suggests that it explores the nucleus using a more compact search strategy than ZLD, in which molecules spend more time trapped in nuclear sub-regions, often retracing their steps^41^. To test this idea, we calculated the distribution of angles between three consecutive displacements in single-molecule trajectories and found a 1.2-fold increase in diffusional anisotropy for ZF5 overall and 1.5-fold inside clusters, compared to ZLD overall (Fig. 3F-H). These changes in anisotropy suggest that Zelda molecules more frequently revisit regions, after its DNA binding activity is perturbed. We hypothesized that this behavior is driven by ZF5 being more frequently trapped in hubs which facilitates its target search^7^, potentially explaining its ability to occupy its new targets despite a reduced residence time.

### IDR interactions stabilize hub lifetimes despite DNA binding deficiencies

We hypothesized that the increase in trapping of ZF5 molecules compared to ZLD could result from a change in the number or temporal stability of hubs. To test this idea, we acquired and analyzed volumetric lattice light-sheet data throughout the duration of nuclear cycles (ncs) 12-14 (Figs. 4A and S6, Movies 5-12). We observed that ZLD forms transient hubs that appear discrete in ncs 12 and 13, and take on a more connected appearance in nc 14. In contrast, ZF5 forms hubs that are longer-lasting than those formed by ZLD but are fewer per nucleus and remain more discrete through ncs 12-14. We found that ZF5 hubs occupy less nuclear volume than ZLD hubs throughout the entire durations of ncs 12-14 (Fig. 4B). To quantify differences in individual hub lifetimes, we performed intensity autocorrelation analysis (Figs. 4C, S7) and found that ZLD, and all mutants we tested, form two populations of hubs - one shorter- and one longer-lived. In nc 14 although ZLD and ZF5 don’t have significantly different proportions of short- and long-lived hub populations, homozygous ZF5 hubs on average persist for seven seconds longer (Figs. 4D-E and Table S6). Strikingly, all mutants we tested exhibit slightly longer average hub-lifetimes than wildtype ZLD with no significant differences in the proportions of short and long-lived hubs with the exception of ΔIDR. Hubs formed by ΔIDR exhibit a severe depletion in the fraction of hubs that are long-lived (~20% for ΔIDR vs. 44-50% for other conditions, Fig. S7C-D), implicating the role of IDRs in stabilizing hubs. As hubs can mediate TF occupancy despite the short residence time of proteins on chromatin by facilitating an increase in binding frequency, we reasoned that the more stable ZF5 hubs could drive its occupation at its new targets by more frequently localizing with them.

**Fig 4.**
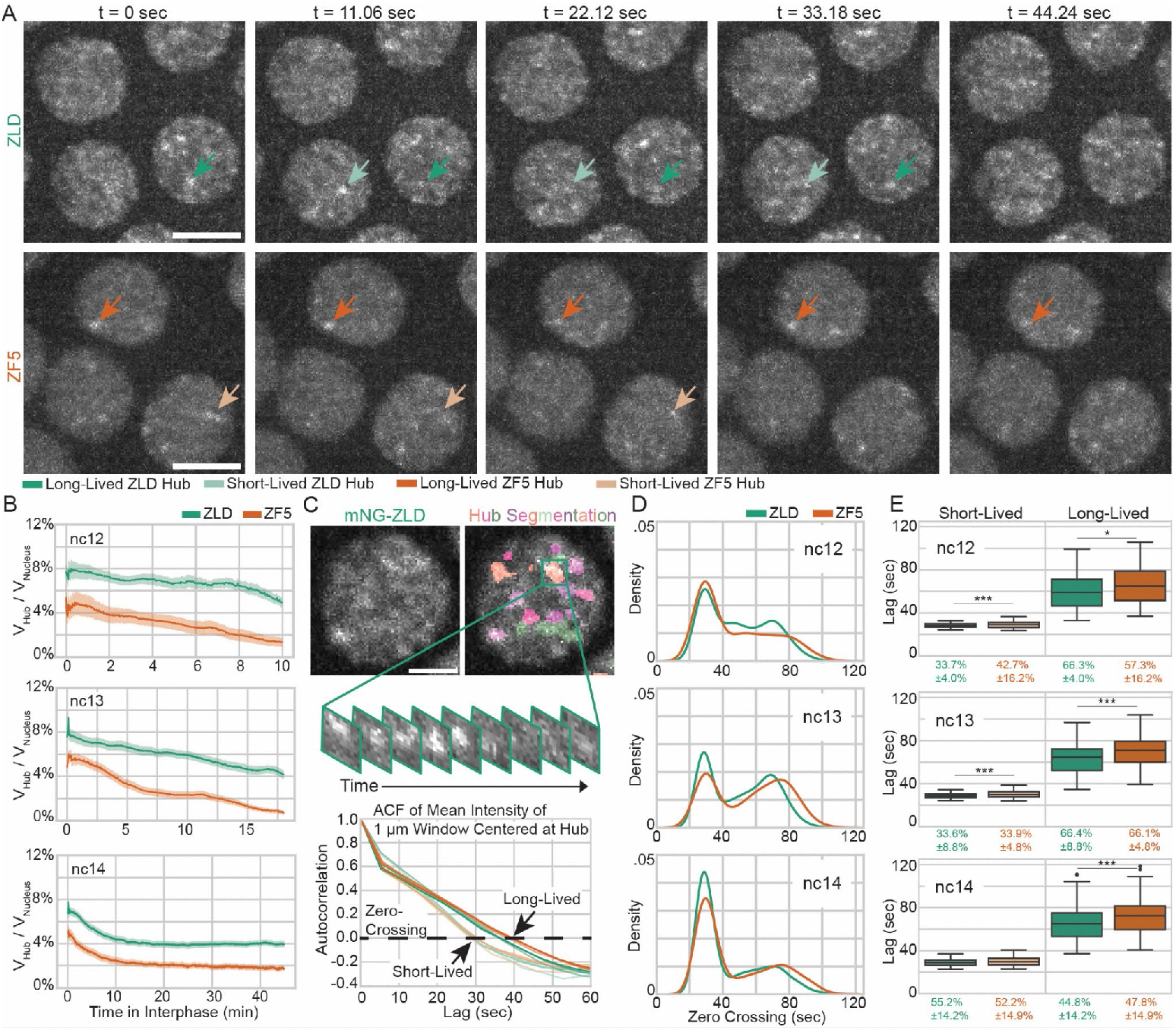
Perturbation of Zelda DBD leads to fewer but longer-lived hubs. **(A)** Images of mNeonGreen-ZLD (top) and mNeonGreen-ZF5 (bottom) over 45 seconds in nuclear cycle (nc) 13. Dark green/orange and light green/orange arrows show representative long-lived and short-lived hubs respectively. Scale bars are 5 microns. **(B)** Fraction of nuclear volume occupied by hubs in interphases of nc12, 13, and 14. For ZLD, 72 nuclei from 4 embryos in nc12, 312 nuclei from 9 embryos in nc13, and 895 nuclei in 13 embryos in nc14, and for ZF5, 341 nuclei from 5 embryos in nc12, 199 nuclei from 6 embryos in nc13, and 846 nuclei from 10 embryos in nc14 were analyzed (statistics summarized in Table S5). **(C)** Hubs were segmented and intensity autocorrelations were calculated in 1 µm^3^ boxes centered on hubs (top). Average autocorrelations for short-lived and long-lived populations in each nc for ZLD (green) and ZF5 (orange) are shown. For nc12, total N is derived from 3 ZLD and 5 ZF5 embryos. For nc13, total N is derived from 6 ZLD and 3 ZF5 embryos. For nc14, total N is derived from 9 ZLD and 9 ZF5 embryos (statistics summarized in Table S6). Scale bar is 2 µm. **(D)** Probability density plots of zero-crossings from autocorrelations. **(E)** Zero-crossings and proportions of short- and long-lived hubs.

### DNA binding deficient Zelda hubs are relocalized to a differentially upregulated gene

To test whether ZF5’s genomic relocalization is driven by a relocalization of hub interactions to its new sites, we performed simultaneous two-color lattice light-sheet imaging to assess hub interactions in the context of nascent transcription at an upregulated target. We selected the gene *Antp* for analysis as it is significantly upregulated and shows increased accessibility in ZF5 embryos (Figs. 2A, S8A). We found that in ZF5 embryos *Antp* is transcriptionally active as early as nc12 and continues to be into late nc14, whereas it is only first expressed in nc14 in wildtype embryos (Movies 13-16). We observed a striking increase in the frequency of hub associations to the transcribing *Antp* locus in ZF5 embryos compared to ZLD embryos (Figs. 5A, S8B-G, Table S7). We quantified the differences in relative TF enrichment^18^ driven by hub interactions at *Antp* and found that there is significant enrichment of ZF5 embryos above the nuclear mean intensity, but not in ZLD embryos (Figs. 5B, S8B-C).

**Fig 5.**
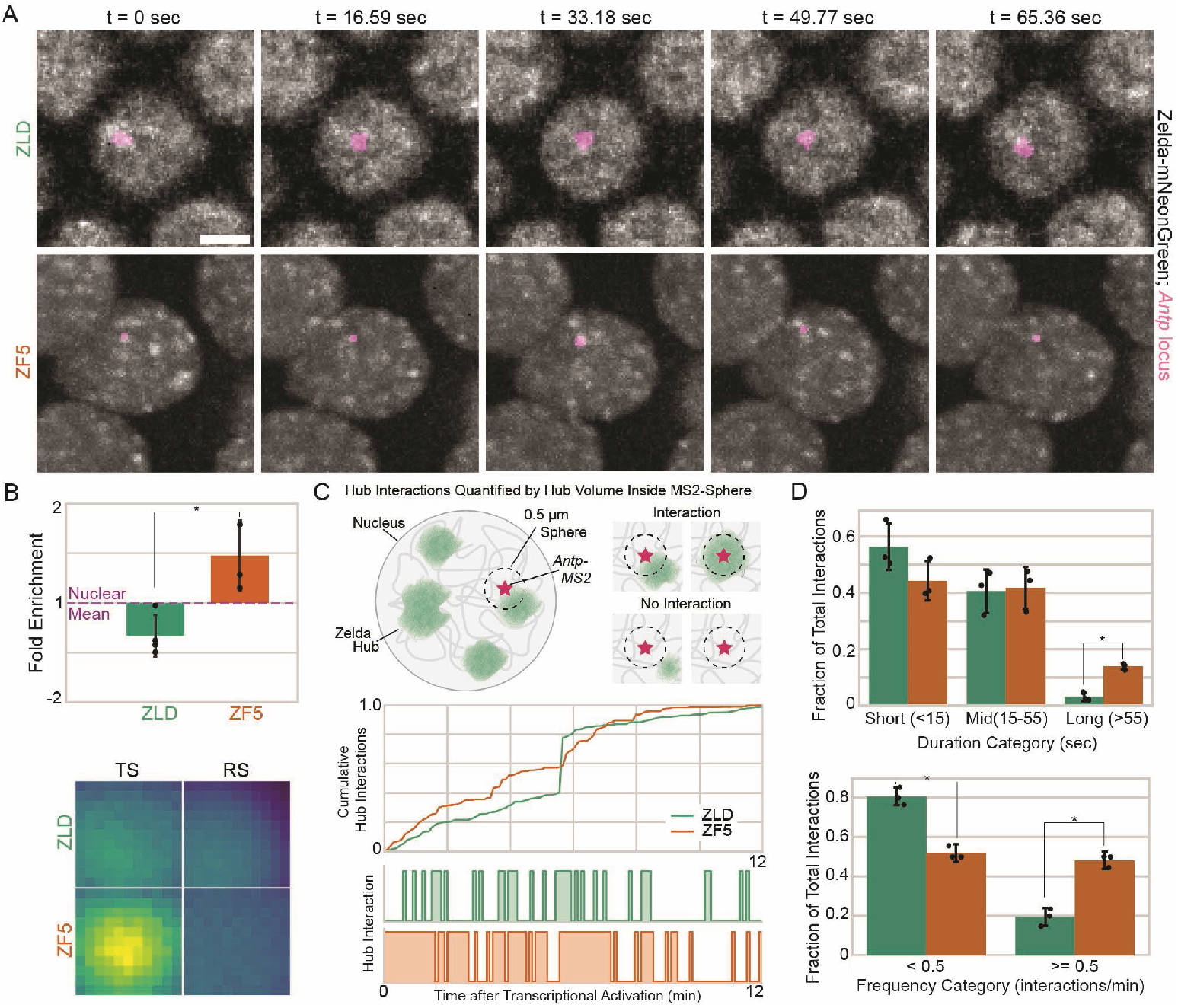
Zelda DBD mutant upregulates new targets through hub relocalization. **(A)** Nuclear distributions (gray) of ZLD (top) and ZF5 (bottom) in nc14 and transcribing *Antp* locus (magenta). Scale bar is 5 µm. **(B)** Fold enrichment of mNeonGreen intensity above the nuclear mean intensity within a 0.55 µm radius centered at *Antp* of ZLD and ZF5. Radial profile plots shown in (Fig. S10). Images show average intensity centered at the transcription site (TS) and random control sites (RS). Images are 1.1 µm x 1.1 µm. **(C)** Hub interactions are quantified using overlapping hub pixels within a 0.5 µm sphere centered at the *Antp*-MS2 spot (TS) or at random control sites (RS). Representative cumulative (top) and individual (bottom) hub interaction traces from one nucleus for ZLD and ZF5 are plotted. **(D)** Fractions of hub interactions per embryo of duration of individual hub interactions (top) and the frequency of interactions (bottom) with the *Antp* locus in nc14 (n=3 embryos for ZLD and n=3 embryos for ZF5, statistics are summarized in Table S7).

We next quantified the kinetics of hub interactions at the *Antp* locus and compared them to randomly selected control spots in the same nucleus in both ZF5 and ZLD embryos (Fig. 5C). We found that the median duration of individual hub interactions increases from ~12 sec in ZLD to ~17 sec in ZF5 embryos, and the frequency of hub interactions is increased from ~0.2/min in ZLD to ~0.5/min in ZF5 embryos (Fig. S8E, Table S7). We noted that there is a large skew towards higher values in the distributions of hub interaction durations and frequencies. We thus categorized the durations of hub interactions into: short (less than 15 seconds), middle (between 15 and 55 seconds) and long (greater than 55 seconds) groups. We found that ZF5 hubs have a significantly greater fraction of long interactions (Fig. 5D, S8D). To assess changes in interaction frequency, we compared groups of less than 0.5 hub interactions per minute and greater or equal to 0.5 hub interactions per minute. We found that ZF5 hubs have significantly more frequent interactions than ZLD hubs at the *Antp* locus (Fig. 5D, S8D). Together, these data reveal that increased occupancy and the differential activity of *Antp* is mediated by an increase in the duration and frequency of ZF5 hub interactions.

### Zelda’s target selectivity is correlated with cofactor binding when its DBD is perturbed

Although Zelda is a key activator of the zygotic genome^34^ and a primary driver of accessibility in the early embryo^32,33^, another pioneer factor, GAGA-factor (GAF), binds independently of Zelda^38,42^. In the absence of Zelda, the morphogenic TF Dorsal has been shown to relocate to sites that are enriched for GAF motifs but depleted in ZLD motifs^43^, and in larval brains where Zelda’s canonical motif alone cannot explain its binding preferences, Zelda peaks are more likely to be found at GAF motifs^44^. We thus hypothesized that the relocalization of ZF5 is driven by interactions with co-factors redistributed to sites that are occupied by GAF in the absence of wildtype Zelda. We compared GAF binding in wildtype and ZF5 embryos and found that GAF binding is highly correlated with sites that are differentially occupied by ZF5, and those that are occupied by both ZF5 and ZLD, but not at sites where ZF5 binding is lost (Fig. 6A-B). GAF binding itself is not altered in the ZF5 background. While, as expected, Zelda motifs are enriched under ZLD peaks, in ZF5, this enrichment is lost and instead an enrichment of GAF motifs is found (Fig. 6C). These peaks are enriched in promoter regions relative to ZLD peaks and also correspond to sites where Dorsal relocates upon Zelda depletion, despite a lack of Dorsal binding motif enrichment (Fig. S9). Strikingly, despite the ZF5 unique sites being occupied by GAF in WT embryos, there is an increase in accessibility at these sites in both ZF5 and ΔZLD embryos. This indicates that the redistribution of co-factor to GAF sites is driving increased accessibility when ZLD activity is lost. These analyses suggest that ZF5’s relocation to new target sites is driven by interactions with co-factors that are themselves redistributed when Zelda’s occupancy at its canonical targets is perturbed. Although this redistribution is correlated with GAF binding and an increase in accessibility in ΔZLD embryos, transcription is further enhanced by the recruitment of ZF5.

**Fig 6.**
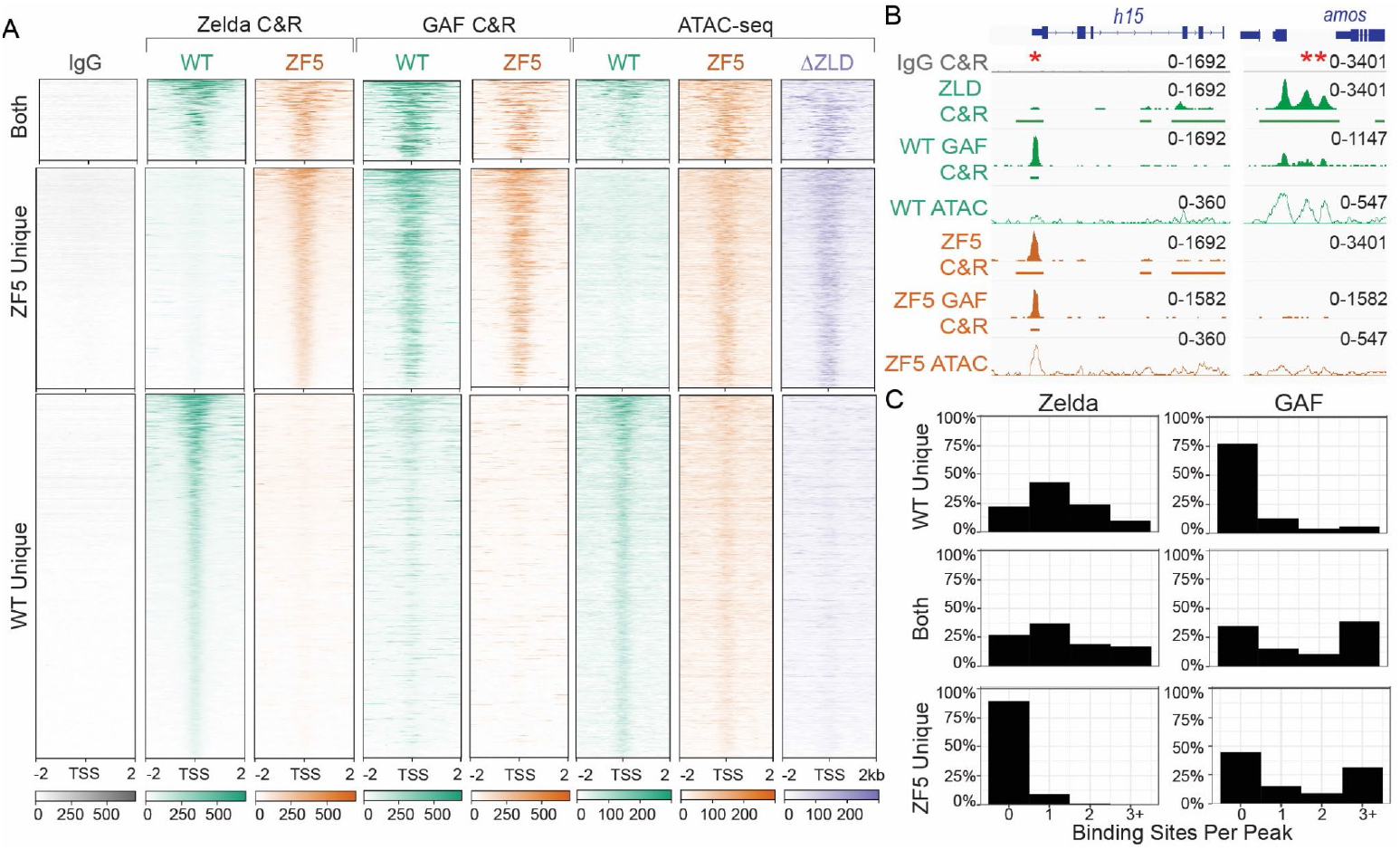
Zelda DNA binding mutant relocates to sites of GAF binding. **(A)** Heatmap of Zelda peaks at TSSs separated by whether they are present in WT only (2268), ZF5 only (1478), or both (653). The heatmaps display ZLD and ZF5 CUT&RUN (C&R), GAF CUT&RUN in WT and ZF5 embryos, and ATAC-seq in WT, ZF5, and ΔZLD embryos. **(B)** Example ZLD, ZF5, and GAF tracks (Solid), ATAC-seq (hollow), and called peaks (solid bars). *denotes a GAF peak where ZF5 is relocalized to. **denotes a ZLD peak that is lost in ZF5. **(C)** Histograms of binding site counts in ZF5s unique binding sites, WT Zelda unique binding sites, and binding sites shared by both. Hits must match the consensus sequence by 90% for Zelda, and 85% for GAF.

## DISCUSSION

The ability to measure molecular kinetics *in vivo* has revealed that TF target site occupancy is not determined by the duration of binding events, but by the frequency of binding, which can be increased by accumulating TFs in the vicinity of their target sites. These local accumulations represent distinct nuclear microenvironments^4^ which are in many cases driven by low-affinity but selective interactions between the IDRs of TFs^21,23^. At super-enhancers such microenvironments (composed of Mediator and RNA Pol II) are more stable in time and exhibit liquid-like droplet properties, and are as such referred to as condensates^9,11,19^. However, persistent condensate-like bodies that last for tens of minutes only represent ~10% of the observed clusters in these studies, with global average lifetimes only lasting on the order of ten seconds^11,14^. The challenges involved in characterizing these ubiquitous transient TF hubs in *in vivo* and endogenous contexts has hindered our ability to understand how hubs are targeted to specific genes and how they then drive TF occupancy at these sites.

Our initial observation that inhibiting the ability of Zelda to recognize its cognate motifs relocalizes it to new target sites (Fig. 2) opened the opportunity to investigate the mechanisms of hub targeting in the context of embryonic development. The fact that the Zelda ZF5 mutant relocalizes to GAF sites along with another co-factor of Zelda, leads us to a model in which hub targeting is dictated by the relative strengths and abundances of co-binding factors and protein-chromatin interactions (Fig. 7). The relocalization of the ZF5 mutant to sites occupied by GAF enforces the idea that site selectivity for most TFs is determined by both their DBD activity and interactions with co-binding factors through non-DBD regions. Many TFs binding sites are low-affinity in nature, and thus depend on the presence of co-binding factors to occupy their sites^45^. Indeed, locally high-concentrations of TFs, such as those found in hubs, can render low-affinity binding sites functional^46^. Given Zelda’s role in mediating the binding of a majority of developmental TFs to their targets^32^, our results suggest that the occupation of a site by a given TF is orchestrated by the recruitment of likely compositionally distinct hubs, dictated by the underlying genomic sequence, which drives the presence of a multitude of binding partners. When Zelda can no longer bind its cognate sites the TFs which Zelda recruits to these sites are now free to relocate and bind to other available targets. ZF5 retains the ability to interact with these co-binding partner TFs and thus relocalizes along with them. The formation and localization of hubs thus represents a complex kinetic balancing act, involving transient low-affinity DNA-protein and protein-protein interactions that determine the recruitment of different transcription factor families to target sites.

**Fig 7.**
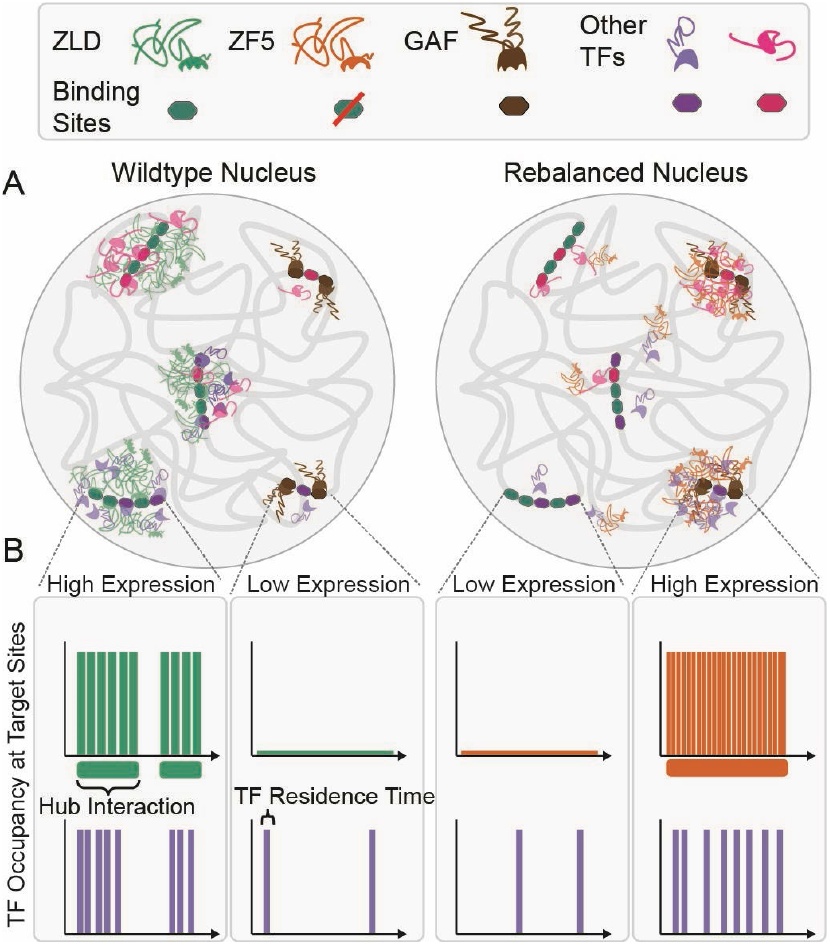
A model for hub relocalization due to a kinetic rebalancing in the nucleus. **(A)** In nuclei from wildtype embryos (left) Zelda binds to its strong DNA motifs, pioneers them and drives transcription by both enabling the binding of and recruiting co-factors and additional TFs in locally enriched hubs. In nuclei from ZF5 embryos (right), Zelda no longer binds its cognate motifs; it and its cofactors are now drawn to orthogonal sites where they form new types of hubs, many of which are bound by GAF. **(B)** Although the residence time of ZF5 on chromatin is severely reduced, its hubs are now more stable in time and interact more frequently with new target sites where they drive high occupancy and expression.

Although it is recognized that condensates and hubs are likely multifactorial they are often viewed as having consistent emergent biophysical properties. This view that key properties such as lifetimes, relative enrichments, and sizes are singular, is based on the widespread assumption of the role of equilibrium thermodynamic processes such as liquid-liquid phase separation in driving their formation^22^. Our finding that the lifetimes of individual Zelda hubs vary considerably within a single nucleus suggests that their stability is tuned in a target-specific manner (Fig. 4). The increase in hub lifetimes at ectopic sites in the ZF5 mutant leads us to speculate that specific co-binding factor interactions are key to tuning the stability and frequency of hub interactions. Our results also highlight potential variability in the functional roles of hubs; for example, hub formation at one target may indeed lead to a loss of occupation at other sites leading to lower gene expression^47^. This sort of kinetic competition may be akin to recent observations on the sequestration of CDK9 in large transcriptional bodies ^48^.

Our observation that the residence times of the ZF5 mutant are reduced almost five-fold over the wildtype protein (Fig. 3) while it forms longer lived hubs that interact more frequently with a target site (Fig. 5) leads to the conclusion that hub stabilities and proximities are key determinants driving TF occupancy at a target. We speculate that underlying combinations of low-affinity binding sites, presence and levels of co-binding factors, epigenetic modifications^49^, and local chromatin organization together tune hub stabilities and interaction frequencies. We posit that these local properties have been evolutionarily optimized to tune TF occupancy at specific loci depending on cellular context. In situations where this fine balance has been disrupted, hubs might be aberrantly stabilized leading to disease^50^. The selection and occupation of gene targets by transcription factors through mechanisms other than specific DNA motif recognition may carry unique advantages at evolutionary timescales as they do not require changing or evolving new DNA binding motifs. We speculate that over the course of evolution such weak but tunable interactions, along with the plastic nature of IDRs, have allowed for maximal changes in phenotype, with minimal perturbations to overall topology of the highly pleiotropic gene regulatory networks that are a hallmark of early development.

While our results here deepen our understanding of the fine balance of interaction kinetics that drive TF hub targeting, further exploration is needed to elucidate the rules that dictate essential properties such as hub composition and formation probabilities, and the roles of non-specific DNA binding vs. IDRs. Interdisciplinary approaches such as we have developed here using a combination of high-resolution volumetric imaging to track hub properties at target loci, single-molecule tracking to understand the impact of hub formation on molecular interactions, and genomics to assess functional effects, will be critical to tackling these fundamental questions on the mechanisms of gene regulation.

## Acknowledgments

We thank Michael Stadler, Michael Eisen, and Melissa Harrison for initial discussions regarding this project and for providing critical fly lines. We thank Xiao-Yong Li in the Eisen lab for providing antibodies against Zelda and fly lines. We thank Alan Boka for preliminary data informing this work. We thank John Lis for providing antibodies against GAF. We thank Daniel Milkie, Srigokul Upadhyayula, Wesley Legant, Tian-Ming Fu, and Eric Betzig for critical support in the construction of the light-sheet microscope used in this work. We thank Ken Zaret, Arjun Raj, and Melike Lakadamyali for helpful discussions regarding this work. We thank all members of the Mir lab for insightful discussion and critical feedback. The work was supported by National Institutes of Health grant DP2HD108775 (to M.M.), Margaret Q Landenberger Foundation (M.M.), Howard Hughes Medical Institute, Freeman Hrabowski Scholars Program (M.M)., and National Science Foundation Graduate Research Fellowship grant DGE-2236662 (to S.F.).

## Author contributions

M.M. conceived and supervised the study. M.M. and A.M. constructed the light-sheet microscope and established single-molecule tracking. A.M. performed and analyzed single-molecule tracking data with the assistance of K.S.. S.F. performed and analyzed volumetric imaging data and wrote software packages. P.R. performed all CUT&RUN experiments with the assistance of Y. H. and with J.Z. performed analysis. P.R. designed and constructed new fly lines and validated them. S.F., A.M., P.R., J.Z., and M.M. interpreted the data, prepared figures, and wrote the initial manuscript. All authors contributed to review and editing.

## Data and materials availability

All the data and code are available as described in the Materials and Methods section. The GEO accession number for genomics data is GSE264096.

## MATERIALS AND METHODS

**Key Resources**

| Reagent or Resource | Source | Identifier | Additional Information |
| --- | --- | --- | --- |
| **Fly Lines** |  |  |  |
| mNG-ZLD | Mir et al. 2018<sup>18</sup> | - | Fly line with an N-terminal mNeonGreen fusion tag inserted at the endogenous Zld locus. |
| mEos3.2-ZLD | Mir et al. 2018<sup>18</sup> | - | Fly line with an N-terminal mEos3.2 fusion tag inserted at the endogenous Zld locus. |
| sfGFP-ZLD | Hamm et al. 2017<sup>36</sup> |  | Fly line with an N-terminal sfGFP fusion tag inserted at the endogenous Zld locus. |
| mNG-ZF5 GLC | This paper | - | Fly line with an N-terminal mNeonGreen fusion tag inserted at the endogenous Zld locus modified with the ZF5 mutation and recombined with a line containing the FRT19A line to generate germline clones for homozygous expression. |
| mEos3.2-ZF5 GLC | This paper | - | As above but with a mEos3.2 N-terminal tag. |
| mNG-ΔIDR GLC | This paper | - | As above but with a deletion of all domains N-terminal of the DNA binding domain and with mNeonGreen N-terminal tag |
| mNG-ZF1-6 GLC | This paper |  | As above but with all 6 Zinc fingers scrambled. |
| Zld<sup>294</sup> GLC | Liang et al., 2008<sup>34</sup> |  | Homozygous Knock-out of Zelda using the FRT19A line described above. |
| His2B-mEos3.2 | Mir et al. 2018<sup>18</sup> | - | Fly line with an H2B-mEos3.2 transgene inserted on chromosome 3. Transgene is expressed ubiquitously under the control of a synthetic tubulin promoter. |
| His2B-EGFP | Mir et al. 2018<sup>18</sup> |  | Fly line with an H2B-eGFP transgene inserted on chromosome 3. Transgene is expressed ubiquitously under the control of a synthetic tubulin promoter. |
| Antp-MS2 | Batut et al. 2022<sup>52</sup> |  |  |
| **Antibodies** |  |  |  |
| Zelda | Eisen Lab (UC Berkeley) | - |  |
| GAF | Lis Lab (Cornell University) |  |  |
| Anti-GFP antibody | Abcam | ab290 |  |
| Anti-beta Tubulin antibody [EPR16774] | Abcam | ab179513 |  |
| H3K4me3 Antibody, SNAP-Certified™ for CUT&RUN | EpiCypher | 13-0041 |  |
| CUTANA™ Rabbit IgG CUT&RUN Negative Control Antibody | EpiCypher | 13-0042 |  |
| **Critical Commercial Reagents** |  |  |  |
| Concanavalin A Magnetic Beads and Activation Buffer | CellSignaling Technology | 93569S |  |
| CUT&RUN pAG-MNase and Spike-In DNAq | CellSignaling Technology | 40366S |  |
| NEBNext® Ultra™ II DNA Library Prep Kit for Illumina® | New England Biolabs Inc | E7645 L |  |
| Agencourt AMPure XP | Beckman Coulter | A63880 |  |
| Protease Inhibitor Cocktail (PIC) | SigmaAldrich | 11697498001 |  |
| NEBNext® Ultra™ II RNA Library Prep Kit for Illumina® | New England Biolabs Inc | E7775 S |  |
| D1000 tapestation assay | Agilent | 5067-5582 |  |
| GlycoBlue™ Coprecipitant | ThermoFisher Scientific | AM9516 |  |
| RNA High Sensitivity tapestation assay | Agilent | 5067-5579 |  |
| Phasemaker™ tubes | ThermoFisher Scientific | A33248 |  |
| Nextera XT DNA Library Preparation Kit | Illumina | FC-131-1024 |  |
| Illumina Tagment DNA Enzyme and Buffer Small Kit | Illumina | 20034197 |  |
| Nextera DNA CD Indexes 24 indexes - 24 samples | Illumina | 20015881 |  |
| TRIzol™ Reagent | ThermoFisher Scientific | 15596026 |  |
| ECL™ Anti-rabbit IgG, Horseradish Peroxidase | Amersham™ | NA934V |  |
| ECL substrate | BioRad | 170-5060 |  |
| Nitrocellulose membrane | BioRad | 1620215 |  |
| Digitonin | Milipore | 300410 |  |
| Spermidine | ThermoFisher | 132740010 |  |
| **Deposited Data** |  |  |  |
| ATAC-seq (WT and ZF5) | This paper | GSE264096 |  |
| RNA-seq (WT and ZF5) | This paper | GSE264096 |  |
| CUT&RUN (ZLD, ZF5, and GAF) | This paper | GSE264096 |  |
| GAF- ChIP-seq | Gaskill et al. 2021<sup>42</sup> | GSE152773 |  |
| Dorsal-ChIP-seq | Sun et al. 2015<sup>43</sup> | GSE65441 |  |
| RNA microarray | Liang et al. 2008<sup>34</sup> | GSE11231 |  |
| **Software** |  |  |  |
| Quot | https://github.com/alecheckert/quot | N/A |  |
| SASPT | <sup>40</sup><br>https://github.com/alecheckert/saspt | N/A |  |
| Custom SMT Analysis | https://gitlab.com/mir-lab/publications/ZeldaAnalysis_2024 and 10.5281/zenodo.13851055 | N/A | Find all code for this in the “single_molecule_analysis” folder. Gitlab has code only. Zenodo includes data CSVs. |
| pyEnRICH | https://gitlab.com/mir-lab/publications/pyEnRICH | N/A |  |
| Custom Hub Analysis, MS2-Hub Interaction Analysis | https://gitlab.com/mir-lab/publications/ZeldaAnalysis_2024 and 10.5281/zenodo.13851055 | N/A | Find all code for this in the “volumetric_image_analysis” folder. Gitlab has code only. Zenodo includes data CSVs and sample images. |
| Imaris Parallel Writer | https://github.com/abcucberkeley/LLSM5DTools | N/A |  |
| micro-SAM | https://github.com/computational-cell-analytics/micro-sam | N/A |  |
| Cellpose 2.0 | https://github.com/MouseLand/cellpose | N/A |  |

### Fly Husbandry

All germline clone embryos were made in similar fashion using the following protocol. mNG-ZF5/+ flies were expanded in large dextrose food base bottles and kept at 25°C. mNG-ZF5/+ virgins were crossed to males ovoD, hs-FLP FRT-19A at 25°C for 48 hour before the adults were flipped and larvae were heat-shocked at 37°C for 3 consecutive days at 1 hour intervals (Fig. S1A). Enclosed mNG-ZF5 adult flies were screened before caged with apple agar plate and fed with yeast paste. Embryos laid overnight from these cages were checked for lethal phenotype for 3 consecutive days before the cages were used. Non heat-shock control bottles were also made to determine ovoD effectiveness or potential non-virgin G0 females. ZF5 mutant embryos fail to gastrulate and begin to show defects in late nuclear cycle 14, consistent with previous observations using complete Zelda knock-out embryos^5,34^.

The mNG-ZF5 FRT-19A mutant files were generated using CRISPR-Cas9 mutagenesis. Briefly, the zld CDS, including the ZF5 mutation, was amplified from ZF5 GLC genomic DNA and then cloned into the existing mNeonGreen-zld HDR plasmid in-frame with the FP and GDGAGLIN linker. For mNG-ΔIDR, the zelda DBD region, consisting of ZF3-ZF6, was amplified using the following primers (Forward: GACCACCCTGCCATCGGG and Reverse: AGCGAGCGGGCAAATGCCGC) and ligated in-frame into the GDGAGLIN linker HDR backbone. Lastly, the mNG-ZF1-6 construct was generated through randomized scrambling of the endogenous ZF amino acid sequence and by utilizing Drosophila-optimized nucleotides. All scrambled ZFs were ligated using multiple fragment In-Fusion ligation into the mNeonGreen-linker HDR backbone. These HDR templates and sgRNA plasmids were sent to Rainbow Transgenic Flies, Inc. (Camarillo, CA) and BestGene Inc. (Chino Hills, CA) and injected into Cas9-expressing embryos. The resulting transgenic adults were recombined onto the FRT-19A chromosome and balanced with the FM7 balancer.

### Western blots

Embryos were collected after incubation for 75 minutes at 25°C. Dechorionated embryos were staged under halocarbon oil 27 and rinsed prior to protein isolation. Around 90-100 embryos were appropriately staged and collected in 1x PBS + PIC. Proteins were extracted under denaturing conditions using a 2x Laemmli SDS sample buffer with 5% BME and heated at 95°C for 5 minutes then chilled as previously described^53^. Samples were centrifuged at 4°C for 10 minutes at 16000 RPM. Supernatant was placed in new 1.5 ml tubes and stored at −20°C until needed. Protein samples were split into two equal volumes and run on 8% acrylamide gel at 200 volts for 5 hours for zelda protein separation and 1 hr for beta-tubulin separation. Proteins were transferred onto nitrocellulose membranes overnight at 4°C at 40 volts and blocked for 45 minutes with 3% milk in 1X TBST (1X TBS, 0.2% Tween) at room temperature. The blots were then incubated overnight at 4°C with either rabbit anti-Zelda or rabbit anti-beta-tubulin (1:1000 dilution) for loading control. Anti-Rabbit HRP antibody was used as a secondary at 1:5000 for 1 hour at room temperature. Blots were treated with ECL substrate and visualized with the BioRad ChemiDoc system (Fig. S1B).

### Embryo collection and mounting for live imaging

Embryos were collected after incubation at 25°C for 75 minutes. Embryos were dislodged from an apple juice-agar collection plate in water, transferred to a cell strainer, and subjected to bleach for 30-45 seconds to remove the chorion. The plate was then rinsed with distilled water until the bleach odor dissipates, embryos were selected with a fine paintbrush and positioned on an agar pad under a dissection microscope. A 25 mm glass coverslip is made adhesive by depositing 20 μl of a solution of double-sided scotch tape dissolved in heptane. Once the heptane has fully evaporated the embryos are transferred to the coverslip by gently tapping.

### CUT&RUN

Embryo collection for each genotype varied slightly due to embryo depositing rate especially for germline clone mutant collections, however typical protocol was completed as followed. Embryos were collected at 25°C for 30 minutes, and incubated for 60 minutes at 25°C. Embryos were dechorionated with 50% bleach for 30-45 seconds with vigorous mixing. Embryos were then rinsed thoroughly and moved to slightly moistened apple agar for staging. A small drop of halocarbon oil 27 was overlaid on the embryos to help staging. Properly staged embryos were collected on a small kimwipe paper before gently rinsed with slow drops of water. The embryos were gently rolled off the kimwipe paper and placed on a double-sided tape and placed on a slide. Several pools of CUT&RUN wash buffer^54^ were placed around the embryos. Using a sterile 18 gauge metal needle, quickly pop embryos while drawing the content into the pools of the wash buffer. Extracted contents were quickly moved in the wash buffer into a 1.5 µl tube and temporarily stored at room temperature while samples were collected. Nuclei were collected by slow centrifugation at room temperature at 1500 RCF for 10 minutes. Nuclei were bound and fragmented using conA beads for 20 minutes at room temperature. Samples were then incubated in primary antibodies at 4°C on a thermomixer with gentle mixing. pAG-MNase was used for fragmentation as directed in manufacturer’s protocols. STOP buffer from ^55^ was used for proper chromatin release. Spike-in DNA was added at 2 pg/ml for normalization. CUT&RUN libraries were made using NEBNext Ultra II NGS library kit and size selected using AMPure XP beads using manufacturer’s protocols. Each CUT&RUN library were verified using a Tapestation and sequenced by NovoGene Co. with pair-end sequencing at 17M reads-depth per library.

### Single-embryo ATAC-seq

Briefly embryos from each genotype were collected for 30 minutes and hand dechorionated after 75 minute incubation at 25°C. Embryos were then mounted on a double-sided tape on a cover slip and overlaid with halocarbon oil 27 for staging. Proper staged ncs 12-13 embryos were removed off double-sided tape and rinsed before they were placed in 1.5 mL tube caps. Embryos were macerated with an 18-gauge needle in lysis buffer (10 mM Tris-HCl pH7.5; 10 mM NaCl, 3 mM MgCl_2_; 0.1% Igepal-630). Samples were quickly spun and placed on dry-ice to snap freeze. Fragmentation and ATAC-seq library amplifications were performed as previously described^56,57^.

### CUT&RUN and ATAC-seq analysis

For ATAC-seq and CUT&RUN, reads were aligned to the Berkeley Drosophila Genome Project (version six) reference genome using bowtie2 with the ‘--sensitive-local’ setting. Samtools was used to convert from .sam to .bam, index, sort, and remove duplicate, unmapped, and multi-mapped reads. For CUT&RUN, peaks were called using macs2 using the ‘-f BAMPE’, ‘-g 129789873’, and ‘--bw 300’ settings using IgG as a control. For ATAC-seq, no IgG was used as a control and --nomodel --nolambda settings were used. Consensus peaks were called using the ‘diffbind’ package in R. Peaks were annotated using the CHIPseeker package and ‘TxDb.Dmelanogaster.UCSC.dm6.ensGene’ annotation. Bigwig files were generated using the Bamcoverage function from the *deeptools* package using the a ‘--binSize 20’, ‘--smoothLength 60’, ‘--centerReads’, ‘--normalizeUsing RPKM’, ‘--effectiveGenomeSize 142573017’. Bigwigs were further normalized by subtracting the chromosomal median from all bins. WT and ZF5 conditions were averaged using bigwigAverage with a ‘-bs 20’. Heatmaps were generated using deepTools computeMatrix and plotHeatmap functions^58^. Profile plots were generated using deeptools plotProfile function.

### Motif analysis

Motif counts within peaks were generated for Zelda, Dorsal, and Gaf. The motif matrices were obtained from JASPAR with the identifiers MA1462.1 (Zld), MA0205.2 (Trl/Gaf), and MA0023.1 (Dl). Peaks called from the Zelda CUT&RUN were sorted into unique to WT, unique to ZF5, ^58^and consensus. Peaks were expanded by 50 base pairs at each end. Motifs falling within these peak regions with a match fraction higher than 0.9 for Dl and Zld and 0.85 for Trl/Gaf were summed for each condition. Histograms of the number of counted motifs per peak were made.

### RNA-seq

Embryos were collected after 30 minutes of laying time and incubated for 75 minutes at 25°C. Embryos were hand dechorionated and mounted onto a double-sided tape and overlaid with halocarbon oil 27. Each embryo was staged to proper nc 12 to nc 13 and then rinsed before they were placed on 1.5 mL microcentrifuge tube caps (3 embryos per cap). Embryos were macerated in 10 µl TRIzol™ using sterile an 18 gauge needle. The caps were then closed and stored on ice until all samples were collected. Samples were briefly spun down and additional Trizol were added. Samples were incubated at room temperature for 5 minutes before equal volume of chloroform was added. This mixture was quickly spun down and moved to Phasemaker™ tubes. Isolation of total RNA was completed as previously described ^59^ with a small modification where GlycoBlue was added according to manufacturer’s instructions as a coprecipitant to help visualize RNA pellets. Isolated total RNA samples were measured for quality using RNA High Sensitivity tapestation assay; samples with RIN 9 or above were used to make final RNA libraries. RNA-seq libraries were made using NEBNext® Ultra™ II RNA Library Prep Kit for Illumina® following manufacturer’s recommended protocol. Libraries’ quality were checked using D1000 tapestation assay. RNA libraries were sequenced by NovoGene Co. using pair-ended sequencing at 27M reads-depth per library.

### RNA-seq analysis

To quantify transcript abundances from RNA-seq data we used the pseudoaligner Kallisto. Reads were aligned to the *Drosophila melanogaster* BDGP6.28 cDNA reference using a pre-built kallisto index^60^. Data was normalized by eliminating transcripts with no reads in more than half of the samples from each condition and then using EdgeR to perform the ‘Trimmed Mean of M-values’ method to correct for compositional differences between libraries. Clear differences between the WT and ZF5 transcript abundances were observed via Principal Component Analysis (Fig. S3). To determine differentially expressed genes, the ‘voom’ method from the ‘Limma’ package in R was applied to the log-transformed counts per million (CPM) data. After fitting linear models to the transformed gene expression data using the lmFit function, we applied the empirical Bayes (eBayes) method to the model fits. Subsequently, we extracted the top DEGs from the eBayes-fitted models. We adjusted for multiple testing to control the false discovery rate using the Benjamini-Hochberg (BH) method. Genes were classified as differentially expressed if they had logFC greater than 1.5 and an adj. p-value less than 0.05.

To analyze the microarray data from Liang et al.^34^, we imported the microarray data using the ‘ReadAffy’ function from the ‘affy’ package to handle .CEL.gz files, which were then normalized employing the Robust Multi-array Average (RMA) method. To identify differentially expressed genes, the ‘limma’ package was used to fit linear models to the expression data. The eBayes method was applied to enhance the statistical power and adjust for multiple testing using the BH method, similar to our RNA-seq data analysis approach. We integrated the CUT&RUN, ATAC-seq, and RNA-seq data sets at promoters and across gene bodies in WT and ZF5 embryos (Table S3). Reads within 2000 bp of TSS of a gene were counted as promoters. Reads across the gene body plus 500 bp at the 5’ and 3’ end were counted as gene body.

### Light-sheet microscope optical paths and configuration

The lattice light-sheet microscope^61^ used in this work is a modified home-built implementation of the instrument similar to the adaptive optics equipped lattice-light sheet^62^ system built based on designs from the Betzig lab at the HHMI Janelia Research Campus. For experiments in this work, the following laser lines were used: 405 nm, 488 nm, and 561 nm. The laser lines were expanded to a diameter of 2 mm combined, passed through a half-wave plate to adjust polarization and relayed into an acousto-optic tunable filter (Quanta-Tech, AA Opto Electronic) to select wavelength and modulate power. The output from the AOTF was then either sent to an optical path to generate a lattice light-sheet excitation pattern or a multi-gaussian beam excitation pattern. For lattice light-sheet generation, the collimated laser beams were expanded along a single dimension using a Powell Lens (Laserline Optics Canada), and the width of the expanded beam was adjusted and collimated using a pair of cylindrical lenses. This stripe of collimated light was relayed onto a grayscale Spatial Light Modulator (SLM) (Meadowlark Optics, AVR Optics) after passing through a second half-wave plate. The diffracted light from the SLM was then relayed onto a custom annular mask to select the minimum and maximum numerical aperture of the light-sheet and block the undiffracted light from the SLM. The light from the annulus was demagnified and projected onto resonant galvanometer (Cambridge Technology, Novanta Photonics) conjugate to the sample plane. The resonant galvanometer was used to mitigate shadowing artifacts and other inhomogeneities in the light-sheet by introducing a slight wobble in the excitation angle. The light is then projected onto a pair of galvanometer scanning mirrors (Cambridge Technology, Novanta Photonics) conjugate to the pupil plane for scanning the light-sheet along x and z optical axes in the excitation coordinate plane. Finally, an excitation objective (Thorlabs, TL20X-MPL) was used to focus the light-sheet onto the sample. The emitted fluorescence was collected by a detection objective oriented orthogonally to the excitation objective (Zeiss, 20×, 1.0 NA), and projected onto a Deformable mirror (ALPAO) positioned in a pupil of the detection path. The light was then split using a dichroic beam splitter (Semrock Di03-R561-t3-25×36) and imaged onto two sCMOS detectors (Hamamatsu ORCA Fusion). The first camera had a green emission filter (Semrock FF03-525/50-25) and a notch filter (Chroma ZET488NF) to reject laser light and the second camera had a red emission filter (Semrock FF01-593/46-25) and a notch filter (Chroma ZET561NF) to reject laser light. The detection path optical aberrations were corrected as previously described by adjusting the deformable mirror (Alpao)^62^.

### Single-molecule imaging using light-sheet microscopy

For single-molecule imaging the gaussian light sheet path was used instead of the lattice light sheet path (see the section on “Light-sheet microscope optical paths and configuration” for more details). Briefly, after passing through the AOTF, the beam was expanded using a Powel lens following which it was relayed onto a custom mask for filtering. The filtered sheet was then projected onto a resonant galvanometer (Cambridge Technology, Novanta Photonics). Finally, an excitation objective (Thorlabs, TL20X-MPL) is used to focus the light-sheet onto the sample. The detection path for the emitted light is the same as described above for the lattice light sheet configuration. It should be noted that for this light path, we bypass the SLM completely in order to ensure higher laser power at the excitation objective.

For all the single-molecule imaging, the 405 nm laser line was kept on constantly during the acquisition period for photoswitching and the 561 nm laser line was used for excitation. Data was acquired at 10 msec and 500 msec exposure times for the fast and slow single-molecule imaging experiments respectively. The excitation laser power was optimized empirically for each exposure time to achieve sufficient contrast for single-molecule tracking and the powers of the photoswitching laser were also optimized empirically to achieve low enough densities of detections to enable tracking. The excitation laser power was 2 mW, 13 mW and switching laser power was 0.2 µW, and 1 µW for 500, and 10 msec exposures, respectively, as measured at the back focal plane of the excitation objective. The same settings were used to acquire control data at each exposure time on His2B-mEos3.2. The number of frames acquired was 8000 and 400 for the 10 msec and 500 msec exposure times respectively corresponding to 80 sec and 200 sec of total imaging time respectively. The acquisition length was optimized such that multiple fields of views could be imaged within each short interphase time while also capturing a sufficient number of trajectories at each position. To optimally position the embryo in the light sheet and to keep track of cell-cycle phase, and nuclear cycle, His2B-eGFP was used.

### Processing of single-molecule tracking data

To process the SMT data we first removed the initial frames during which the fluorophores were bleached. Then we took maximum projections of the data to obtain our nuclear mask. Localization and tracking of the single molecule trajectories was performed using the open source software package *quot*^*63*^. For the 10 msec data the following key parameters on *quot* were used: (i) for detection -method = “identity”, Spot detection settings = “llr”, k = 1.3, w = 15, t = 20, (ii) for localization - method = “ls_int_gaussian”, window size = 9, sigma = 1.0, ridge = 0.0001, maximum iterations = 10, damp = 0.3, camera gain = 1, camera background = 108 (iii) for tracking - method = “euclidean”, pixel size (in µm) = 0.108, search radius = 0.7 µm, maximum blinking = 0 frames. For the 500 msec data, the following parameters in *quot* were used: (i) for detection - method = “identity”, Spot detection settings = “llr”, k = 1.3, w = 15, t = 23, (ii) for localization - method = “ls_int_gaussian”, window size = 9, sigma = 1.0, ridge = 0.0001, maximum iterations = 10, damp = 0.3, camera gain = 1, camera background = 108 (iii) for tracking - method = “euclidean”, pixel size (in µm) = 0.108, search radius = 0.3 µm, maximum blinking = 0 frames. Once we obtained our list of tracked molecules from *quot* we filtered them using the nuclear mask as described earlier to ensure we analyzed only trajectories that were occurring inside the nucleus.

### Quantification of residence times

Residence times were calculated from single-molecule data acquired at 500 msec exposure times as described previously^18,64^. Briefly, imaging at 500 msec exposures effectively blurs out molecules that are not still for a significant portion of the exposure allowing us to selectively localize and track the bound and slowly diffusing population of molecules. The longer exposure times also allow us to use lower excitation laser powers, thus limited photobleaching and providing longer trajectories. To estimate the genome average residence time, survival probability curves were calculated by accumulating data from each experimental condition at each nuclear cycle. An objective threshold of two seconds^18^ was applied to the minimum trajectory length and probabilities below 10^−3^ were not considered for fitting. These filters are applied to remove the effects of tracking errors and slowly diffusing molecules. The survival probability curves were then fit to a double exponential model of the form F *exp(-k_ns_ *t) + (1 - F) * exp(-k_s_ * t) using the curve_fit function in Python, where k_s_ is the slower off-rate and k_ns_ is the faster/nonspecific off-rate. An exponential weighting function was used to ensure proper estimation of the slower off-rate. The inferred off-rate k_s_ is biased by photobleaching and chromatin motion so bias correction was performed as previously described: k_s,true_=k_s_-k_bias_, where k_bias_ is the slower-off rate estimate from fitting the survival probabilities from His2B data. The genome wide residence-times were then calculated as 1/k_s,true_. Errors reported were estimated from the standard-deviations of the fit-parameters using standard propagation of error methods.

### Analysis of fast single-molecule tracking data

To analyze the 10 msec single molecule trajectories to infer the diffusion kinetics we used a variational Bayesian method called State Array based Single Particle Tracking^40^. This method does not a priori assume a specific number of diffusive states. Instead, it processes all the recorded trajectories and selects a model that comprehensively describes the trajectories with the minimum combination of state parameters (diffusion coefficient and localization error) as possible. Consequently, it produces average posterior state occupancies for a state array evaluated on the experimental trajectories. For the model selection, it considers a range of diffusion coefficients from 0.01 µm^2^/s to 100 µm^2^/s representing the range of physiologically relevant diffusion coefficients reported in the literature. This package is publicly available at https://saspt.readthedocs.io/en/latest/. For our analysis, we used the following specific parameters in the SASPT program:

Likelihood type: RBME

Pixel size: 0.108 (1 pixel = 0.108 µm)

Frame interval: 0.0125433 ms

Concentration parameters: 1

Max iterations: 200

Split Size: maximum trajectory length

Sample Size: number of trajectories

Using SASPT we were able to generate the diffusion coefficient occupancy plots for each protein category both overall and inside the clusters. Furthermore, in order to assign individual diffusion coefficients to each trajectory for downstream analysis, we first weighted the range of diffusion coefficients by their corresponding occupancies for each trajectory. We calculated the geometric mean of these weighted coefficients which was then assigned to each trajectory. Then we segregated the trajectories into three kinetic bins defined as follows: bound trajectories (DC≤0.08 um^2^s^−1^) based on the inflection point from the His2B diffusion coefficient occupancy plot, intermediate trajectories (0.08 um^2^s^−1^<DC<0.5 um^2^s^−1^), and fast trajectories (DC≥0.5 um^2^s^−1^) based on the inflection point of the ZLD diffusion coefficient occupancy plot (Fig. S5D).

To determine the clustering behavior from the single molecule trajectories, we first calculated the average position of the trajectories by considering the mean x and y locations. Average trajectory positions were considered because we wanted to quantify the diffusion kinetics inside the clusters and hence needed to assign diffusion coefficients to individual trajectories. We then used density-based spatial clustering of applications with noise (DBSCAN) to determine the location of the clusters and the trajectories that comprised the cluster. For the DBSCAN analysis, we specified that there had to be a minimum of 10 points (“min. samples”) to define a cluster and the maximum distance between any two points in the cluster was 0.2 µm (“max. distance”). These values were selected by initially plotting the number of clusters as a function of the max. distance to determine where the plot showed an elbow, i.e. after what threshold value of max. distance the number of clusters did not change significantly, a common analysis technique used in DBSCAN to determine the appropriate parameters. The parameters chosen were further validated by qualitatively comparing the DBSCAN predictions to the raw spatial maps of the single molecule trajectories. These values were kept constant between the two protein categories.

For the anisotropy analysis, we first filtered the data sets to remove all the bound trajectories since they can bias the angle distribution. We then calculated the angle between all consecutive jumps while excluding all jumps that were less than 0.2 µm based on the jump distribution of His2B (Fig. S5G). We then calculated the fold-anisotropy metric as the probability of observing a backward jump (with an angle between jumps in the range [180° - 30°, 180° + 30°]), divided by the probability of observing a forward jump (with an angle between jumps in the range [0° - 30°, 0° + 30°]).

Single molecule trajectories were taken from 8 different embryos for ZLD and 3 different embryos for ZF5. For calculating the standard deviation for the bound fraction, fold-anisotropy, and average diffusion coefficient, we used bootstrapping to sample 50% of the data 20 individual times. In order to calculate statistical significance where relevant, we used the Mann-Whitney test since our data was not normally distributed as confirmed by the Kolmogorov-Smirnov test. Statistics are summarized in Table S4.

### Volumetric imaging using lattice light-sheet microscopy

Volumetric imaging used for live imaging of Zelda or MCP used a multi-bessel lattice sheet with maximum numerical aperture to minimum numerical aperture ratio of 0.4/0.3 488 nm (used to excite mNeonGreen and sfGFP) and 561 nm (used to excite mCherry). We performed two types of imaging (1) a higher laser power short duration imaging (over two to four minutes) just to assess hub morphology with increased spatial resolution using a higher 488 nm laser power (0.580 µW) and (2) a lower laser power long duration imaging (one to two hour duration) to assess whole cell cycles and lifetimes using laser powers of 0.226 µW 488nm and 1.44 µW 561 nm. Two color channels were acquired simultaneously for a volume of 18.9 µm sampling z-planes spaced 0.3 µm apart with an exposure time of 30 msec. Each volume was acquired every 3 seconds during acquisition for a total time of ~5.53 seconds in between stacks.

### Analysis of volumetric imaging data to quantify hub properties

All volumetric imaging was pre-processed prior to downstream analysis pipelines by first conducting GPU-accelerated 3D image deconvolution using CUDA^65^. All datasets were deconvolved using input PSF taken by bead images on the LLSM and using 5 iterations for Richardson-Lucy based deconvolution. We used a custom Imaris converter leveraging fast [Tiff or Zarr] file readers to generate Imaris files for data visualization and rendering^66^. After deconvolution, images were subjected to nuclear segmentation. To segment nuclei in our dataset, we created a custom model using Cellpose 2.0^67^. Ground truth data was generated using a mixture of micro-Sam^68^ a napari plugin for “segment anything”^69^ and manual correction on wildtype Zelda images. These data were then used to train Cellpose 2.0. The resulting model was then used to segment all slices of each acquired dataset individually. A custom post-processing pipeline was then utilized in order to stitch the individual slices back together and to interpolate any slices of called objects that were missed in segmentation. We then implemented a nearest neighbor algorithm to track nuclei over the course of interphase in each nuclear cycle.

To quantify hub properties, we created a custom analysis pipeline. Nuclei were first normalized to their mean intensity to assess local enrichments of TFs above nuclear background. This pipeline segments hubs by using a median filter to remove noise followed by image erosion and reconstruction. The reconstructed image was subtracted from the median filtered image to first create a binary mask of high density regions. Then, we called local maxima peaks to be used as markers for watershed segmentation in order to separate hubs that might be fused together in the binary mask since we often see hubs as less discrete and more amorphous. We then used region props to quantify different properties of the hubs such as integrated intensity, mean intensity, and size. We then filtered these hubs by their mean enrichment. Using a cumulative distribution function we saw that wild type Zelda hubs reach 0.5 at an enrichment score about 1.4 times the nuclear mean intensity. We used this cut off as a means to clear any hub that might have a minimal and non-significant enrichment compared to the nuclear mean intensity (Fig. S6C).

To quantify the lifetime of hubs, we used the data acquired after the first 20 frames (~2 min) and before the last 20 frames of each nuclear cycle to avoid confounding effects of chromatin reorganization during early interphase and early prophase. This timeframe is then segmented into a series of 20 frame intervals. At the beginning of each interval, each hub in each nuclei is localized and a 1 µm^3^ box is centered around the hub (Fig. 4C). The center of the box window is adjusted to account for the movement of the nucleus. The mean signal within this box is used to calculate autocorrelations. These autocorrelations were then smoothed using a savgol filter with a window length of 7 and poly order of 2. The zero-crossings were interpolated from the resulting curves and used to measure the hub lifetimes. We verified that lifetimes and hub calling were independent from differences in expression and signal to noise in various mutant backgrounds and embryo to embryo by comparing the hub enrichment vs. resulting zero-crossing and observed no clear correlation between the two measurements (Fig. S6A).

Each condition was randomly subsampled to 100 hubs per condition for statistical testing. Significance was measured using Kruskal-Wallis test followed by Mann-Whitney U-Test for pairwise comparisons. The proportion of all hubs within short-or long-lived groupings are reported below the box-plot along with the standard deviation between embryos replicates. N-values and statistics are summarized in Table S5 for hub properties and S6 for hub lifetimes.

### Analysis of volumetric imaging data to quantify hub enrichment at MS2 sites

Preprocessing was completed as stated in the above section for hub analysis. To then quantify the average enrichment of transcription factors and nascent transcription, we created the pyEnRICH package to calculate radial enrichment around the site of transcription to estimate increased local concentration compared to nuclear background as done previously^18^. This script uses a difference-of-gaussians filter followed by a percentile threshold in order to identify spots of nascent transcription. Then we find non-overlapping random spots in the nucleus to compare to the transcription site. By taking 1.1 µm x 1.1 µm windows we are able to average across time and populations of nuclei in order to estimate the average enrichment using a radial profile of the transcription factor channel centered at the spot of nascent transcription. We remove MS2 sites at the edges of the nucleus to avoid improper results stemming from the boundary of a bright nucleus to low background levels in the cytoplasm. We also included a random nuclear spot control to compare the MS2 site to to assess if hub interactions are random or specific to the locus. To identify a random spot, we orient a 1.5 µm radius from the center of the MS2 site in a random direction and then ensure this new spot and a radius of 0.6 µm is located fully within the nucleus. We find three non-overlapping random spots per nucleus and average the resulting enrichment from these spots and compare with the MS2 spot.

After using pyEnRICH, we created an additional custom pipeline that interpolates the center MS2 position for any dropped/non-segmented points for all time points after onset of nascent transcription up until the last time point per tracked nucleus segmented. Using these updated positions, we then quantified hub interactions with the MS2 spot and with random sites. Since Zelda hubs, like most *Drosophila* transcription factors, form amorphous, transient, that are not completely discrete, we found that pairwise distance of hub centroid and MS2 centroid coordinates were insufficient to describe interactions. Instead, hub interactions were called based on overlapping segmented hub masks and a 0.5 µm radius sphere centered at either the MS2 or random spot (Fig. 4C). Hubs must be overlapped in the sphere for at least 2 frames (~11 seconds) to be considered as interacting with the locus as opposed to due to random fluctuations. The duration (length of consecutive time points containing overlapped pixels) and the frequency (amount of hub interactions called in one nucleus over the total time of transcriptional activity) were then quantified and plotted. Additionally, random spots are identified as described above. We assess the hub-interactions between three random spots per nucleus and plot alongside hub-interactions with the MS2 site in order to differentiate between random and specific effects.

A Mann-Whitney U Test was performed on all pairwise relationships in the duration and frequency plots. N-values and statistics are summarized in Table S7.

## Movie Captions

**Movie 1. Volumetric Imaging of all Zelda mutants in nc13**. 4D volumetric imaging data showing side-by-side comparison of mNeonGreen-ZLD, mNeonGreen-ZF5/+, mNeonGreen-ZF5, mNeonGreen-ΔIDR/+, and mNeonGreen-ZF1-6/+ during interphase of nc13. Scale bar = 5 µm. (Related to Fig. 1).

**Movie 2. Single-Molecule Imaging for determining residence times**. Single molecule data acquired at an exposure time of 500 milliseconds for mEos3.2-His2B, mEos3.2-ZLD, and mEos3.2-ZF5. (Related to Fig. 3).

**Movie 3. Single-Molecule Imaging for quantifying diffusion kinetics of mEos3.2-ZLD**. Single molecule data acquired at an exposure time of 10 milliseconds for mEos3.2-ZLD (Related to Fig. 3).

**Movie 4. Single-Molecule Imaging for quantifying diffusion kinetics of mEos3.2-ZF5**. Single molecule data acquired at an exposure time of 10 milliseconds for meos3.2-ZF5. (Related to Fig. 3).

**Movie 5. Volumetric Imaging of ZLD and ZF5 in nc14**. 4D volumetric imaging data showing side-by-side comparison of mNeonGreen-ZLD and mNeonGreen-ZF5 during interphase of nc14 pre-cellularization. (Related to Fig. 4).

**Movie 6. Volumetric Imaging of ZLD**. 4D volumetric imaging data showing mNeonGreen-ZLD through nc12-14. (Related to Fig. 4).

**Movie 7. Volumetric Imaging of ZF5**. 4D volumetric imaging data showing mNeonGreen-ZF5 through nc12-14. (Related to Fig. 4).

**Movie 8. Volumetric Imaging of sfGFP-ZLD**. 4D volumetric imaging data showing sfGFP-ZF5 in nc14. (Related to Fig. 4).

**Movie 9. Hub Segmentation**. Example of 4D hub segmentation for mNeonGreen-ZLD. Zelda-mNeonGreen is shown in gray, and hub masks are shown in fire LUT (purple/red) (Related to Fig. 1 and 4).

**Movie 10. Volumetric Imaging of ZF5/+**. 4D volumetric imaging data showing mNeonGreen-ZF5/+ through nc12-14. (Related to Fig. 4).

**Movie 11. Volumetric Imaging of ΔIDR/+**. 4D volumetric imaging data showing mNeonGreen-ΔIDR/+ through nc12-14. (Related to Fig. 4).

**Movie 12. Volumetric Imaging of ZF1-6/+**. 4D volumetric imaging data showing mNeonGreen-ZF1-6/+ through nc12-14. (Related to Fig. 4).

**Movie 13. Simultaneous volumetric Imaging of ZLD in the context of nascent transcription of *Antp***. Volumetric imaging data of mNeonGreen-ZLD (gray) and MCP-mCherry (magenta) localized to *Antp-MS2* in late nc14. Cytoplasmic background of MCP-mCherry is due to the protein lacking a NLS to maximize signal to noise within nuclei. (Related to Fig. 5).

**Movie 14. Simultaneous volumetric Imaging of ZF5 in the context of nascent transcription of *Antp* in nc12**. Volumetric imaging data of mNeonGreen-ZF5 (gray) and MCP-mCherry (magenta) localized to *Antp-MS2* in nc12. (Related to Fig. 5).

**Movie 15. Simultaneous volumetric Imaging of ZF5 in the context of nascent transcription of *Antp* in nc13**. Volumetric imaging data of mNeonGreen-ZF5 (gray) and MCP-mCherry (magenta) localized to *Antp-MS2* in nc13. (Related to Fig. 5).

**Movie 16. Simultaneous volumetric Imaging of ZF5 in the context of nascent transcription of *Antp* in nc14**. Volumetric imaging data of mNeonGreen-ZF5 (gray) and MCP-mCherry (magenta) localized to *Antp-MS2* in nc14. (Related to Fig. 5).

