## Supplementary Information for "A fine kinetic balance of interactions directs transcription factor hubs to genes"

[1]Department of Cell and Developmental Biology, Perelman School of Medicine, University of Pennsylvania; Philadelphia, PA 19104, USA.
[2]Center for Computational and Genomic Medicine, Children's Hospital of Philadelphia; Philadelphia, PA 19104, USA.
[3]Developmental, Stem Cell, and Regenerative Biology Graduate Group, Perelman School of Medicine; Philadelphia, PA 19104, USA.
[4]Howard Hughes Medical Institute, Children's Hospital of Philadelphia; Philadelphia, PA 19104, USA.
[5]Roy and Diana Vagelos Program in Life Sciences and Management, University of Pennsylvania; Philadelphia, PA 19104, USA.
[6]Epigenetics Institute, University of Pennsylvania Perelman School of Medicine; Philadelphia, PA 19104, USA.

### List of Supplementary Materials

**Fig. S1:** Creation of mutant Zelda fly lines.
**Fig. S2:** Characterization of hubs
**Fig. S3:** CUT&RUN and RNA-seq in ZLD and ZF5 embryos
**Fig. S4:** Differential chromatin accessibility
**Fig. S5:** Single molecule tracking data analysis
**Fig. S6:** Analysis pipeline for Zelda hub stability across mutants
**Fig. S7:** Hub lifetimes quantified across Zelda mutants.
**Fig. S8:** ZF5 accumulation at Antp.
**Fig. S9:** Zelda mutant relocalization.

**Movie 1.** Volumetric Imaging of all Zelda mutants in nc13.
**Movie 2.** Single-Molecule Imaging for determining residence times.
**Movie 3.** Single-Molecule Imaging for quantifying diffusion kinetics of mEos3.2-ZLD.
**Movie 4.** Single-Molecule Imaging for quantifying diffusion kinetics of mEos3.2-ZF5.
**Movie 5.** Volumetric Imaging of ZLD and ZF5 in nc14.
**Movie 6.** Volumetric Imaging of ZLD.
**Movie 7.** Volumetric Imaging of ZF5.
**Movie 8.** Volumetric Imaging of sfGFP-ZLD.
**Movie 9.** Hub Segmentation.
**Movie 10.** Volumetric Imaging of ZF5/+.
**Movie 11.** Volumetric Imaging of ΔIDR/+.
**Movie 12.** Volumetric Imaging of ZF1-6/+.
**Movie 13.** Simultaneous volumetric Imaging of ZLD in the context of nascent transcription of Antp.
**Movie 14.** Simultaneous volumetric Imaging of ZF5 in the context of nascent transcription of Antp in nc12.
**Movie 15.** Simultaneous volumetric Imaging of ZF5 in the context of nascent transcription of Antp in nc13.
**Movie 16.** Simultaneous volumetric Imaging of ZF5 in the context of nascent transcription of Antp in nc14.

**Table S1.** Summary of RNA-seq experiments and associated file names submitted to GEO
**Table S2.** Summary of CUT&RUN and ATAC-seq experiments and associated file names submitted to GEO
**Table S3.** Summary statistics of differentially expressed genes (download .csv from supplementary materials)

**Table S4.** Summary statistics for single-molecule data.
**Table S5.** Summary statistics of hub properties from Fig. 1 and 4
**Table S6.** Summary statistics of hub lifetimes from Fig. 4
**Table S7.** Summary statistics of hub-target gene interactions from Fig. 5

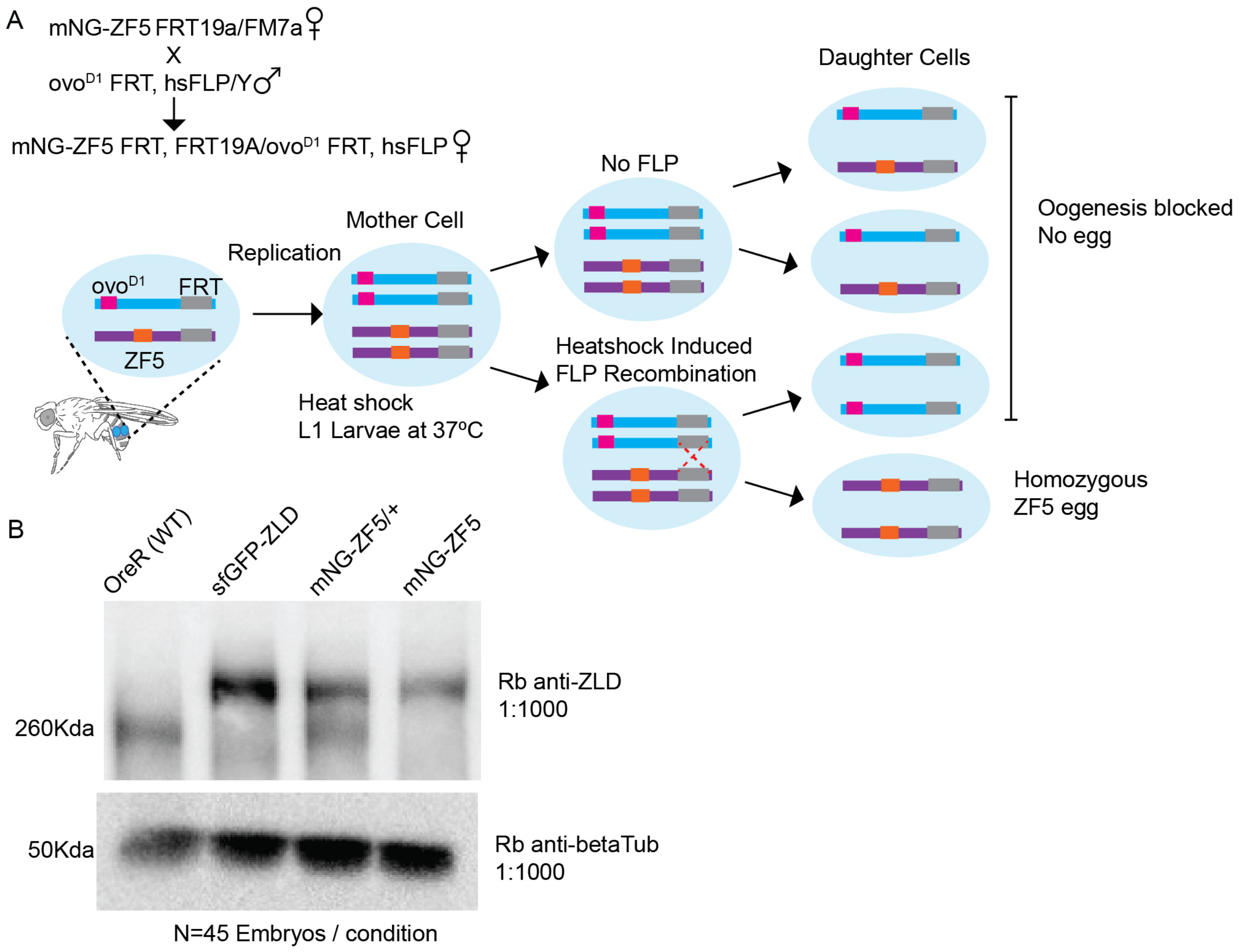


**Fig S1: Creation of mutant Zelda fly lines. (A)** Scheme for generation of germline clones containing homozygous mutations for Zelda using the FLP/FRT system (figure adapted from https://bdsc.indiana.edu/stocks/recombinases/dfs.html). **(B)** Western blots to assess if any wild type Zelda protein is present in embryos generated from germline clones.

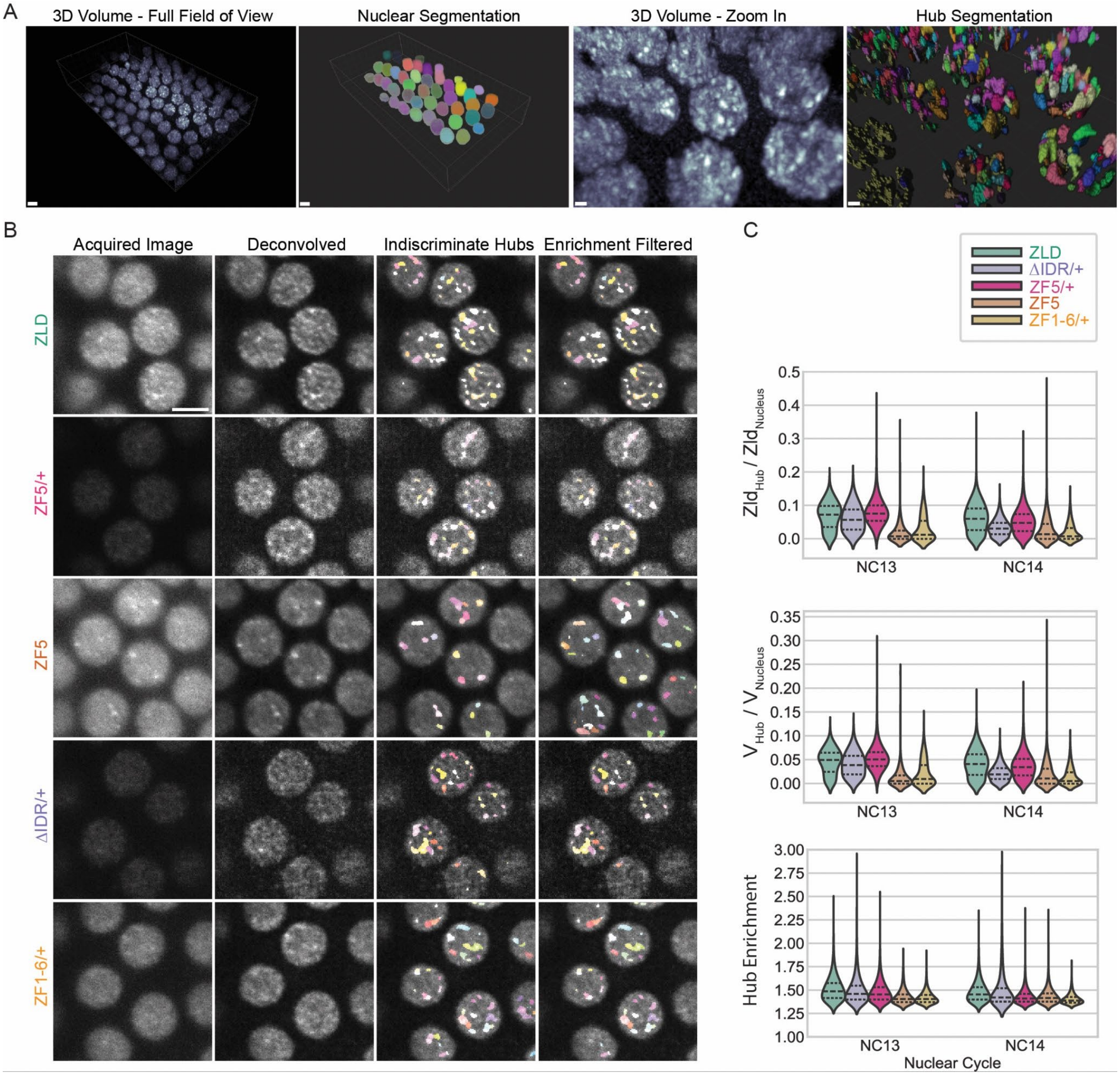


**Fig S2: Characterization of Zelda hubs. (A)** Representation of hub analysis: volumes of a layer of nuclei on the *Drosophila* embryo surface are acquired during imaging and deconvolved (left). Scale bar = 5 µm. Nuclei are segmented in 3D using a trained machine learning model (middle left). Scale bar is 5 µm. Zelda shows non-uniform distributions within single nuclei (middle right). Scale bar = 1 µm. Hub segmentation is performed in 3D using a custom analysis pipeline (right). Scale bar = 1 µm. **(B)** High contrast imaging of mNeonGreen tagged Zelda embryos (ZLD, ZF5/+, ZF5, ΔIDR/+, or ZF1-6/+) in the middle of nc13 before and custom hub segmentation pipeline outputs. Images shown are from a single z-slice in each volume. Acquired images column is shown with consistent contrast adjustment to assess expression levels, while the second column is shown with individual contrast limits adjusted to better visualize hubs. Third column shows initial segmentation output where hubs are indiscriminately called, and third column is after filtering when only hubs with enrichment scores at 0.5 cumulative probability of the wildtype ZLD are considered. Scale bar = 5 µm. **(C)** Quantification of biophysical hub properties including proportion of Zelda within hubs (sum intensity) compared to all nuclear Zelda (sum intensity), the volume fraction of hubs within the nucleus, and hub enrichment (mean hub intensity normalized using mean nuclear intensity) from a snapshots taking in the middle of interphase during nc 13 and 14. N-values and statistics are summarized in Table S5.

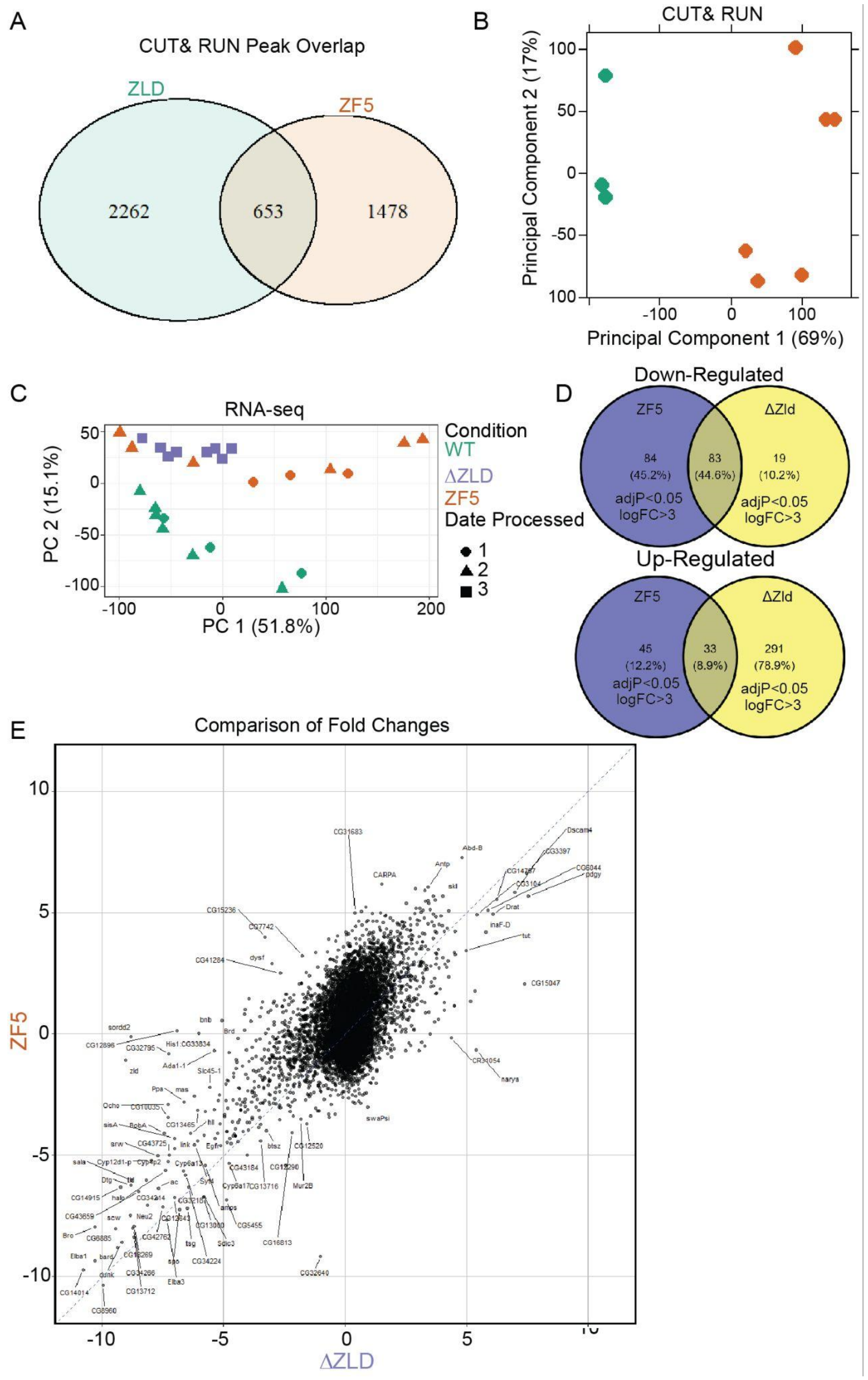


**Fig. S3. CUT&RUN and RNA-seq (A)** A comparison of overlap in identified peaks in CUT&RUN experiments on Wildtype and ZF5 embryos. **(B)** Principal component analysis of replicates from which peaks were calculated in (A). **(C)** Principal component analysis on RNA-seq data. **(D)** Overlap of differentially regulated genes in ZF5, and Zelda knockout embryos **(E)** Scatter plot of all ZF5 vs. Zelda knockout RNA-seq gene expression data.

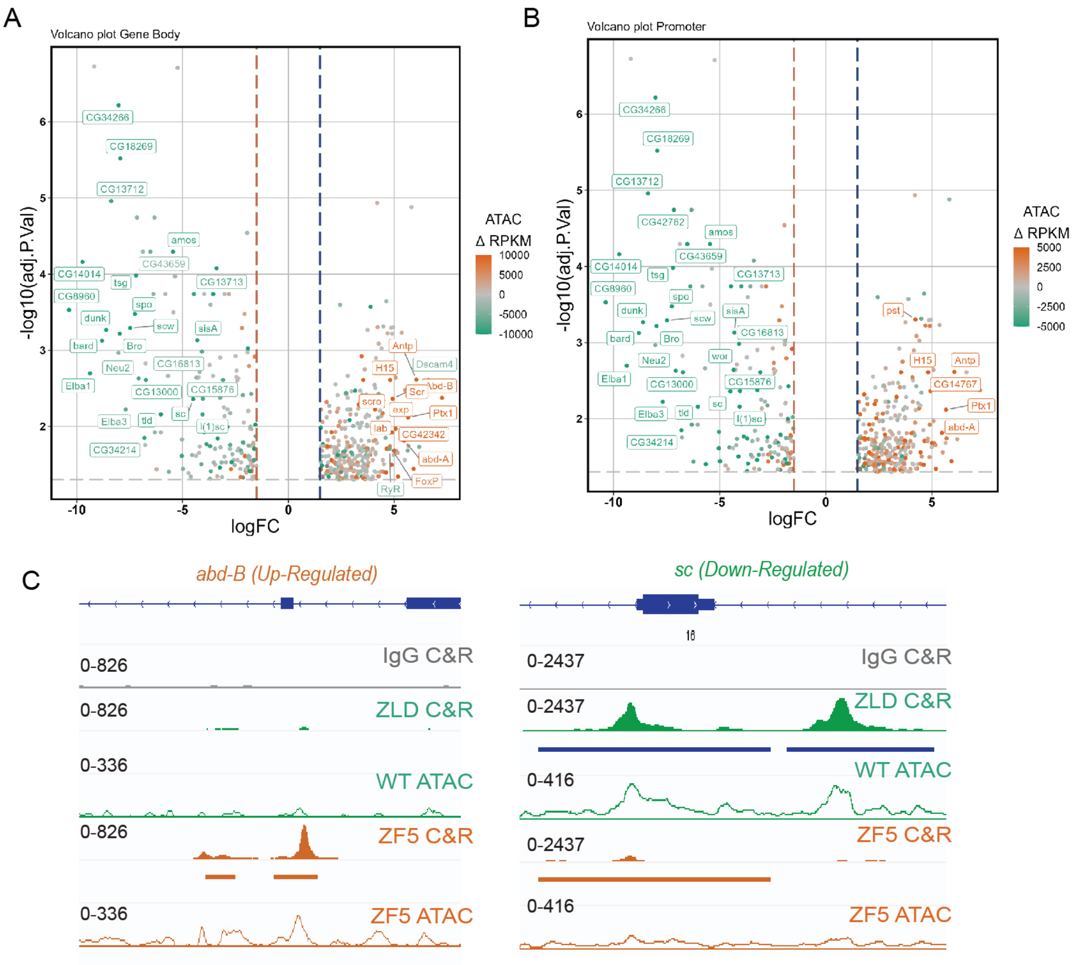


**Fig. S4. Changes in accessibility at up- and down-regulated genes (A)** Volcano plots of genes up- and down-regulated in ZF5 mutant measured via RNA-seq. Genes with a significance <0.01 and Log Fold Change > 4 and a difference in RPKM > 5000 (from CUT&RUN) are labeled . The gray dotted line marks adjust P-value = 0.05, blue and red dotted lines are +/-1.5 LCF. Points are colored by the difference in Zelda RPKM between WT and ZF5 mutants across the gene body of each gene. **(B)** Points are colored by the difference in ATAC RPKM between WT and ZF5 mutants within 2kb of the TSS. **(C)** Example ZLD and ZF5 CUT&RUN tracks (solid), ATAC (hollow), and called peaks (solid bars below tracks) of up- and down-regulated genes.

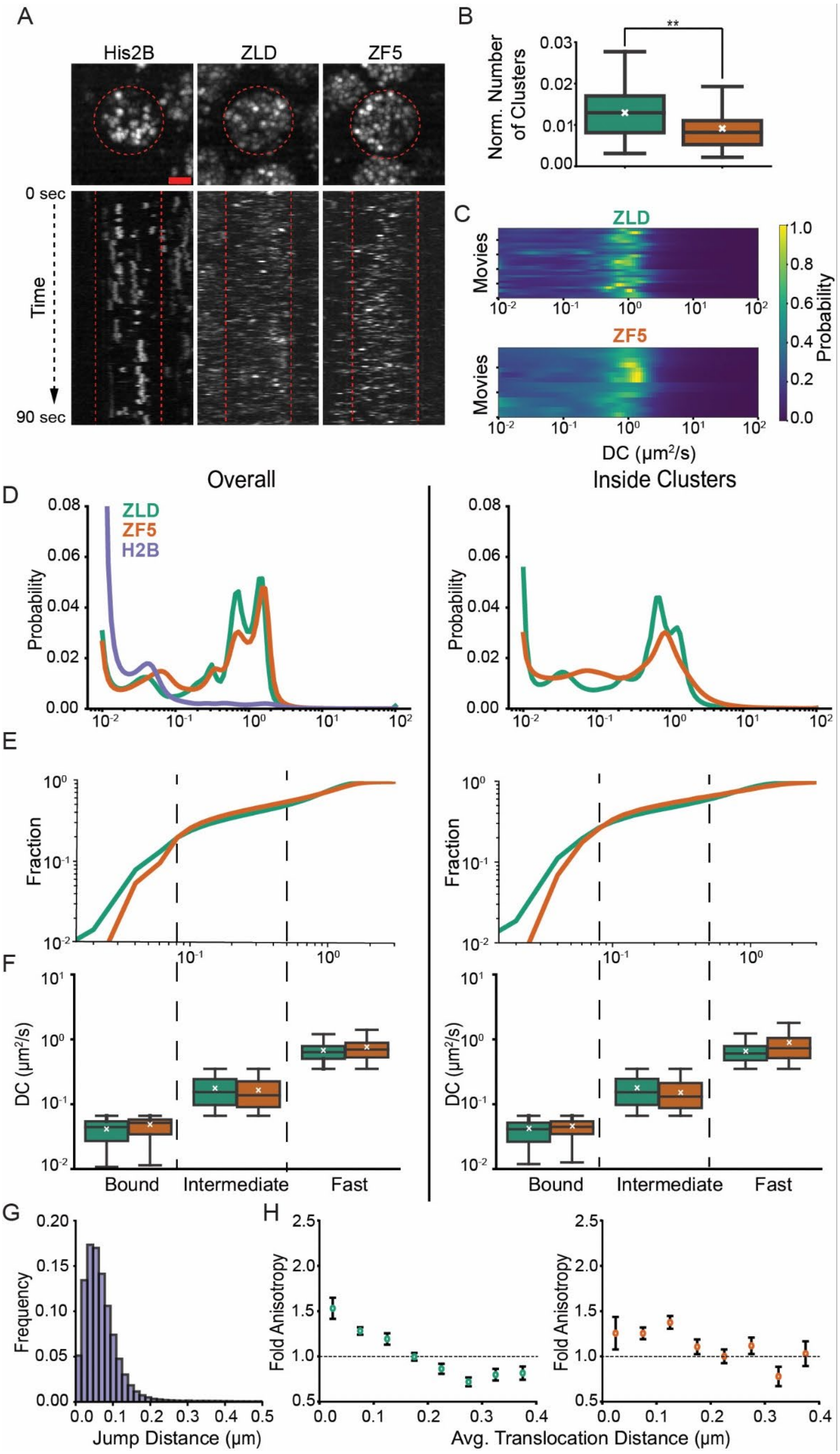
A
His2B
ZLD
ZF5
0 sec
Time
90 sec
B
Norm. Number of Clusters
**
C
ZLD
ZF5
Movies
DC (μm²/s)
Probability
D
Overall
Inside Clusters
ZLD
ZF5
H2B
Probability
E
Fraction
F
DC (μm²/s)
Bound
Intermediate
Fast
G
Frequency
Jump Distance (μm)
H
Fold Anisotropy
Avg. Translocation Distance (μm)

**Fig. S5. Single molecule tracking data analysis**. **(A)** Maximum intensity projections over 90 seconds of raw single molecule tracking data. Red circles indicate nuclear boundaries. Kymographs (x-t maximum projections) of single molecule detections of mEos3.2-ZLD, mEos3.2-ZF5, and H2B-mEos3.2. Acquired with an exposure time of 500 msec, scale bar is 2 μm. **(B)** Quantification of cluster numbers normalized by the total trajectories per nucleus for ZLD (n=172) and ZF5. (n=84). **(C)** Heatmaps of the probability spectra of diffusion coefficients obtained for each movie (one movie/row). Data was analyzed over 23 movies from 7 embryos for ZLD and 14 movies from 3 embryos for ZF5. **(D)** Diffusion coefficient spectra for ZLD, ZF5 and H2B for all trajectories (left) and for ZLD and ZF5 trajectories inside clusters (right). A total of 70,827, 40,307, and 51,158 tracks were measured for ZLD, ZF5 and H2B cases respectively overall, and 14,516 tracks and 3,375 tracks were measured for ZLD and ZF5 respectively in clusters **(E)** Cumulative distribution of diffusion coefficients (DC) for ZLD and ZF5 overall (left) and inside clusters (right). Black dashed lines are cutoffs for bound, intermediate, and fast trajectories. **(F)** Quantification of the average diffusion coefficient in each kinetic bin for ZLD and ZF5 overall (left) and inside clusters (right). White cross is the mean, error bars represent standard deviations from bootstrapping analysis. **(G)** Histogram of jump distances from H2B trajectories used to determine the distance threshold for anisotropy analysis. **(H)** Fold anisotropy as a function of average translocation distance (over two consecutive jumps) for ZLD (left) and ZF5 (right) overall. Error bars represent standard deviation from bootstrapping analysis. Dashed line indicates an isotropic distribution.

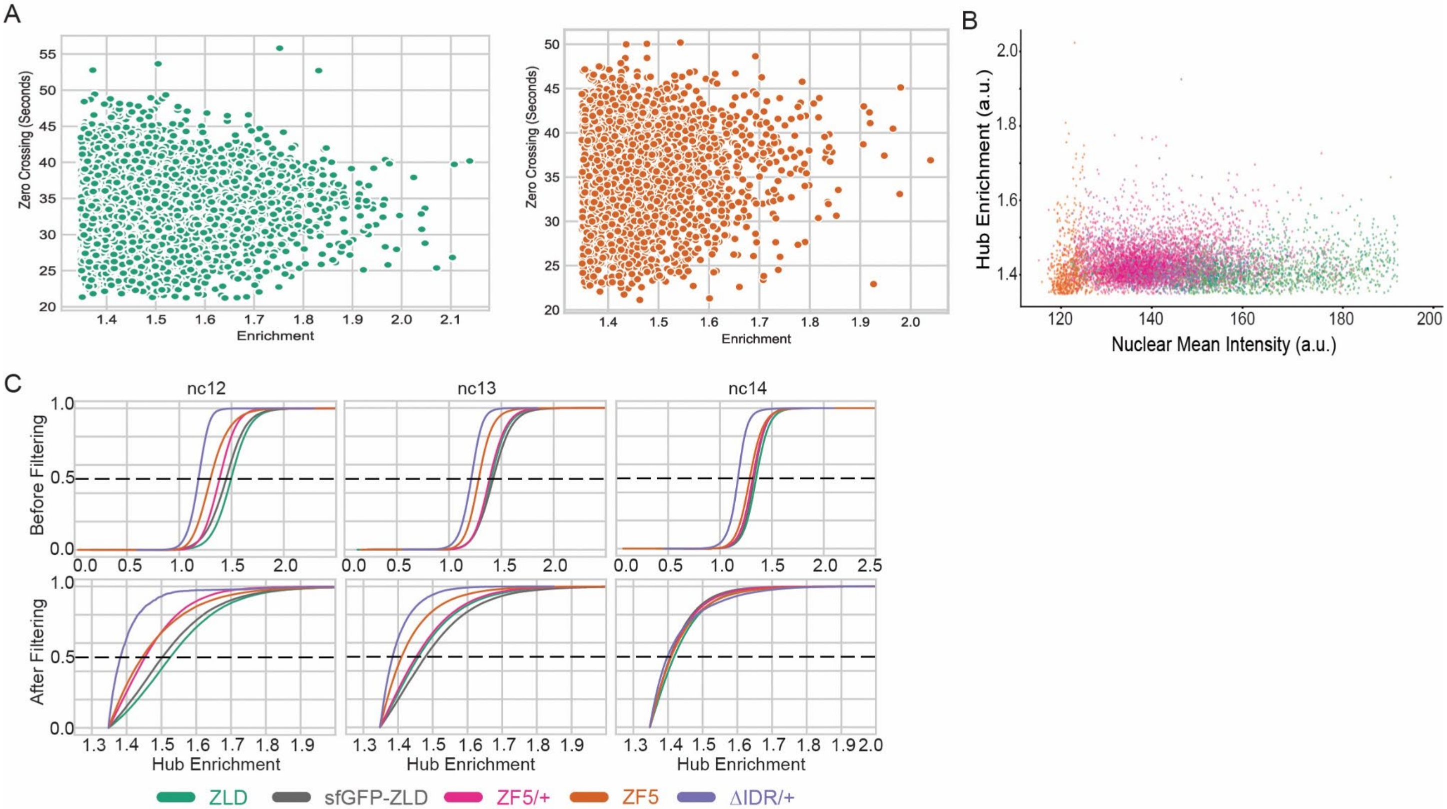


**Fig. S6: Analysis of Zelda hubs across mutants. (A)** Quantification of hub enrichment along with the zero crossing in seconds representing the hub lifetimes show no correlation in either ZLD (top) or ZF5 (bottom) between enrichment and lifetime of each hub. **(B)** Scatter plot showing hub enrichment as a function of nuclear mean intensity. Hub enrichment quantification is not impacted by signal-to-noise difference among mutants. **(C)** Cumulative distributions of hub enrichment for each Zelda mutant before and after filtering (top and bottom respectively).

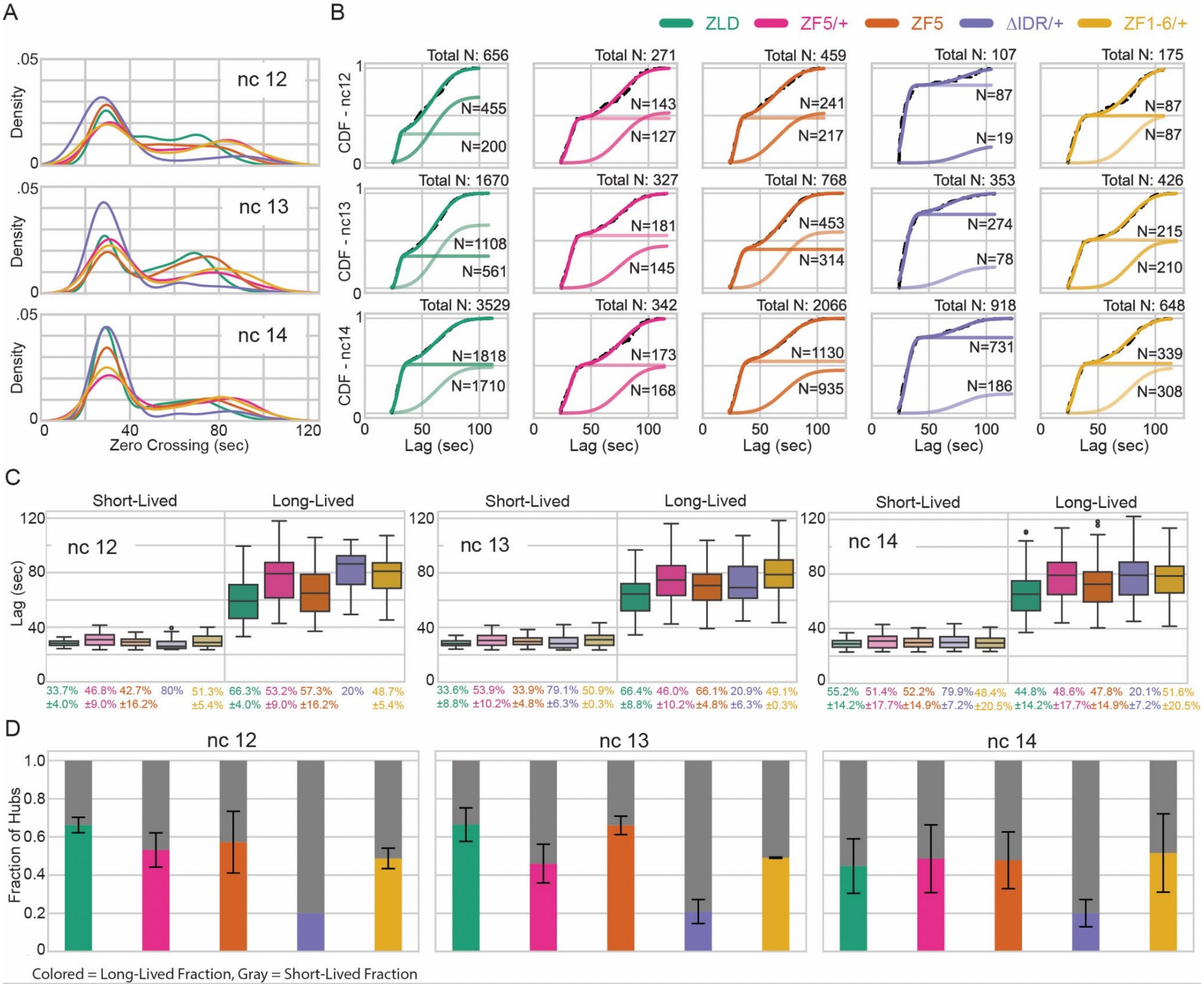

**Fig. S7: Hub lifetimes quantified across Zelda mutants. (A)** Density plots for Zelda mutants showing zero crossing of the autocorrelation of intensity of a 1 μm$^3$ box centered at each hub. Distributions are not normal but suggest a two-component mixture of hubs. **(B)** Cumulative distributions of zero crossings from original data (black dotted line), fitted CDF (solid dark color), and two gaussian components (lighter colors). N values provided for the number of hubs fit into these components. For nc12, total N is derived from 3 ZLD, 4 ZF5/+, 5 ZF5, 2 ΔIDR/+, and 3 ZF1-6/+ embryos. For nc13, total N is derived from 6 ZLD, 3 ZF5/+, 3 ZF5, 3 ΔIDR/+, and 3 ZF1-6/+ embryos. For nc14, total N is derived from 9 ZLD, 3 ZF5/+, 9 ZF5, 4 ΔIDR/+, and 4 ZF1-6/+ embryos. **(C)** Zero-crossing distributions shown in box-plots from each component (short- and long-lived) for nc12, nc13, and nc14 for Zelda mutants. Proportion of hubs within each of the conditions is as written underneath each box plot **(D)** Proportion of hubs in each life-time category. A Kruskal-Wallis test followed by Mann-Whitney U Test was performed among all pairings in the same component in the same nc. All mutants are significantly different from ZLD. Other pairings and all statistical analyses are summarized in Table S6.

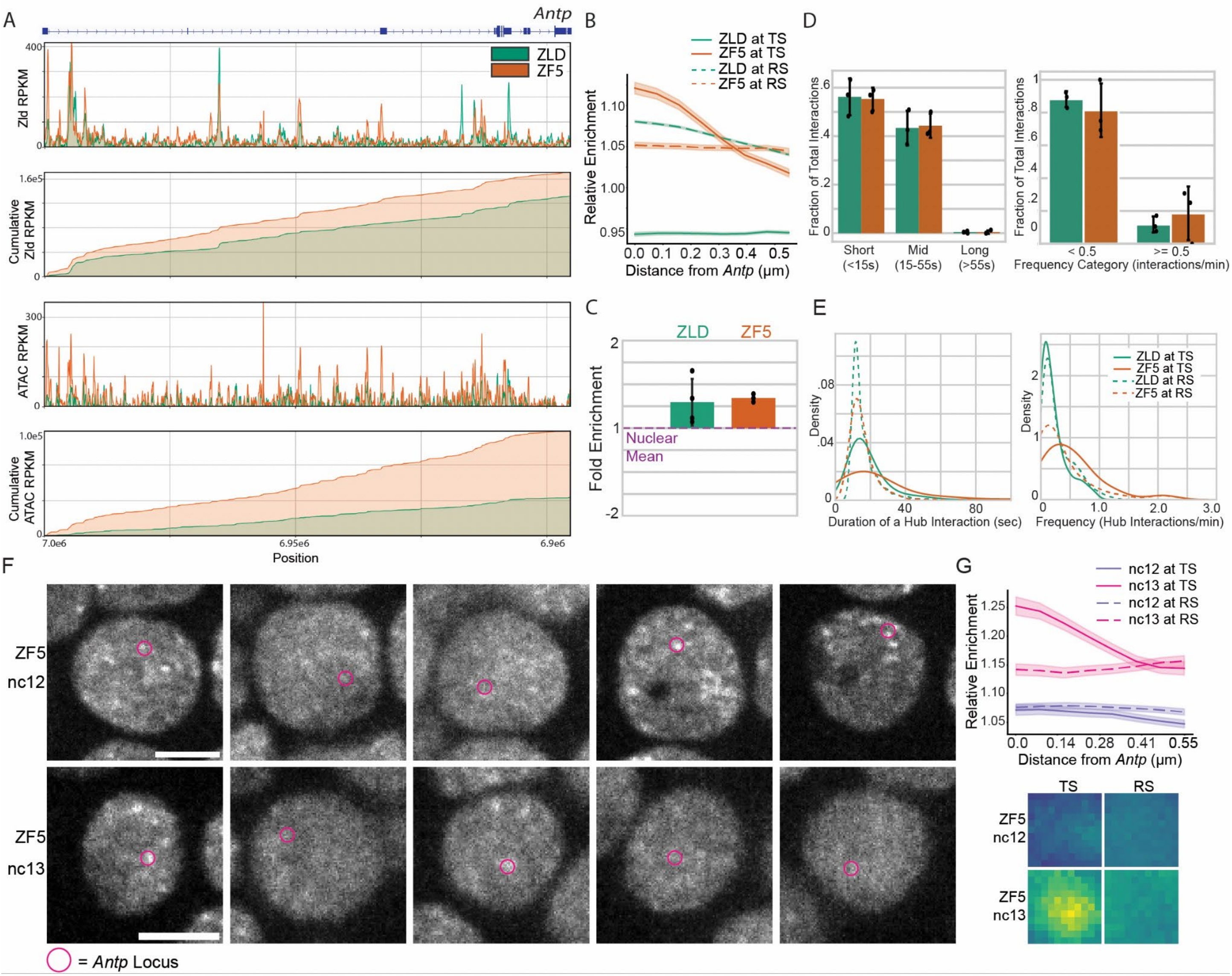


**Fig. S8. ZF5 accumulation at *Antp*.** **(A)** Measured and Cumulative RPKM from CUT&RUN against ZLD and GAF, and ATAC-seq, in WT and ZF5 embryos, along the *AntP* locus. **(B)** Average radial profile with standard error centered at the *Antp* shows ZLD or ZF5 enrichment as a function of distance from MS2 spot center (or transcription site (TS)) or at a control spot randomly placed in the nucleus (or random spot (RS)). **(C)** Fold enrichment of mNeonGreen intensity above the nuclear mean intensity within a 0.55 μm radius centered at a random spot within ZLD and ZF5 nuclei (n=3 embryos for ZLD and n=3 embryos for ZF5, more statistics summarized in Table S7). **(D)** Fractions of hub interactions per embryo of duration of individual hub interactions (left) and the frequency of interactions (right) of a random spot within ZLD and ZF5 nuclei in nc14. **(E)** Distribution of the frequency and duration of hub interactions at *Antp* locus (TS) or random spot (RS) (n=172 nuclei for ZLD and n=65 nuclei for ZF5, more statistics summarized in Table S7). **(F)** Representative images (single slice in Z) of five different nuclei for ZLD and ZF5. Zelda is shown in grayscale while the location of the MCP-mCherry at *Antp-MS2* locus is shown as a pink circle on the images. Scale bar is 5 μm. **(G)** Cropped images show average intensity centered either at the transcription site (TS) or random site (RS) in the nucleus. Image size is 1.1 μm x 1.1 μm. All cropped images have the same contrast adjustment. Average radial profile with standard error centered at the *Antp* shows Zelda ZF5 in either nc12 (purple) or nc13 (pink) enrichment as a function of distance from MS2 spot center. Two embryos were analyzed in nc12 and one in nc 13.

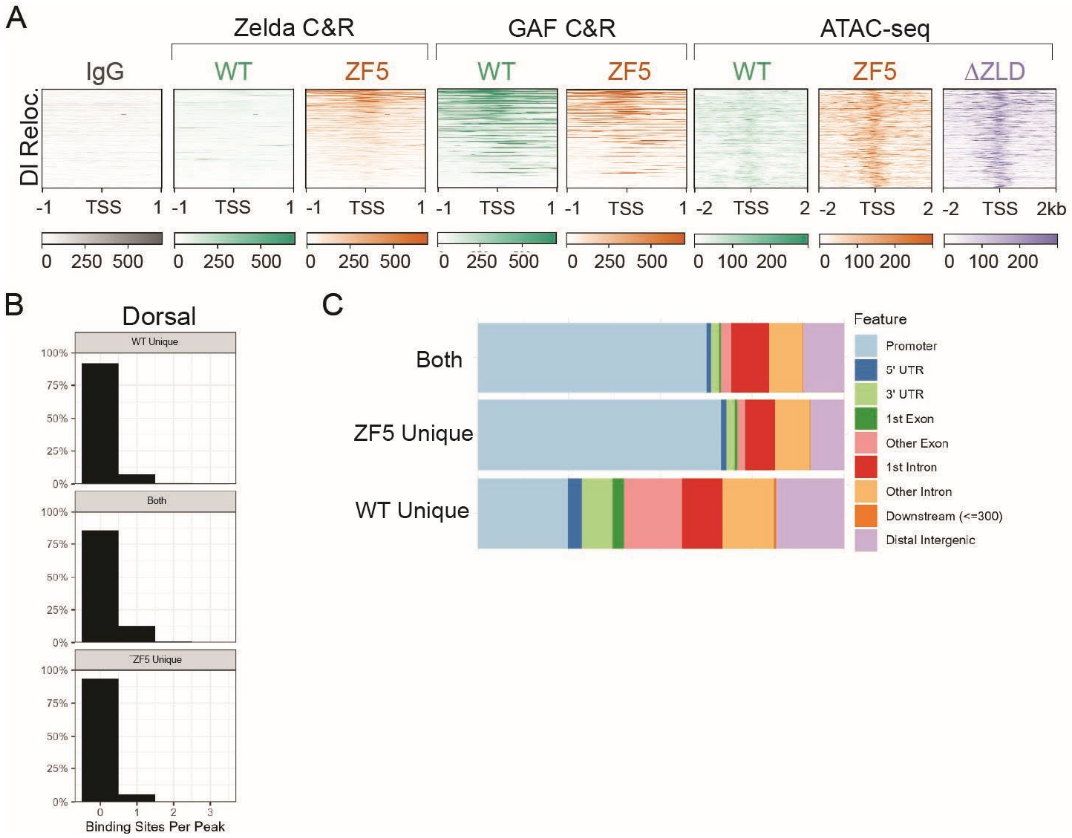


**Fig. S9: Zelda DNA binding mutant relocalization. (A)** ZLD and GAF CUT&RUN (C&R), and ATAC-seq in WT and ZF5 at sites of Dorsal relocalization in ΔZLD null embryos as identified in Sun et al. 2015[43]. **(B)** Histograms of Dorsal binding motif counts in ZF5s unique, WT Zelda unique, and shared binding sites. **(C)** Annotation of peak call locations.

**Table S1. Summary of RNA-seq experiments and associated file names submitted to GEO**

| Sample/Filename | Condition | Day | Genotype | Read Count |
|---|---|---|---|---|
| H2B_GFPwt1_1_1 | WT | 2 | H2B-eGFP (III) | 21044634 |
| H2B_GFPwt2_2_1 | WT | 2 | H2B-eGFP (III) | 17390264 |
| H2B_GFPwt2_3_1 | WT | 2 | H2B-eGFP (III) | 17693844 |
| H2B_GFPwt2_5_1 | WT | 2 | H2B-eGFP (III) | 23905163 |
| H2B_GFPwt3_1 | WT | 2 | H2B-eGFP (III) | 16472166 |
| H2B_GFPwt4_1 | WT | 2 | H2B-eGFP (III) | 21418487 |
| znf5mtGLC2_1_1 | Zf5 | 2 | mNGzld_znf5mt_FRT19A/mNGzld_znf5mt_FRT19A | 13801043 |
| znf5mtGLC2_6_1 | Zf5 | 2 | mNGzld_znf5mt_FRT19A/mNGzld_znf5mt_FRT19A | 20463584 |
| znf5mtGLC2_8_1 | Zf5 | 2 | mNGzld_znf5mt_FRT19A/mNGzld_znf5mt_FRT19A | 13055948 |
| znf5mtGLC2_9_1 | Zf5 | 2 | mNGzld_znf5mt_FRT19A/mNGzld_znf5mt_FRT19A | 15760098 |
| znf5mtGLC2_12_1 | Zf5 | 2 | mNGzld_znf5mt_FRT19A/mNGzld_znf5mt_FRT19A | 29243336 |
| znf5mtGLC2_13_1 | Zf5 | 1 | mNGzld_znf5mt_FRT19A/mNGzld_znf5mt_FRT19A | 20783123 |
| mNGznf5_GLC1_1 | Zf5 | 1 | mNGzld_znf5mt_FRT19A/mNGzld_znf5mt_FRT19A | 19518017 |
| mNGznf5_GLC | Zf5 | 1 | mNGzld_znf5mt_FRT19A/mNGzld_znf5mt_ | 18432907 |

| | | | | |
|---|---|---|---|---|
| 2_1 | | | FRT19A | |
| mNGznf5_GLC3_1 | Zf5 | 1 | mNGzld_znf5mt_FRT19A/mNGzld_znf5mt_FRT19A | 23425471 |
| sfGFPzldwt_1B_1 | WT | 1 | sfGFPzld-wt FD1 | 21583776 |
| sfGFPzldwt_3_1 | WT | 1 | sfGFPzld-wt FD1 | 13681957 |
| sfGFPzldwt_5B_1 | WT | 1 | sfGFPzld-wt FD1 | 23554319 |
| Zld294_GLC_1_1 | Zld294 | 3 | Zld294_FRT19A/Zld294_FRT19A | 20494620 |
| Zld294_GLC_2_1 | Zld294 | 3 | Zld294_FRT19A/Zld294_FRT19A | 21489357 |
| Zld294_GLC_3_1 | Zld294 | 3 | Zld294_FRT19A/Zld294_FRT19A | 21032008 |
| Zld294_GLC_4_1 | Zld294 | 3 | Zld294_FRT19A/Zld294_FRT19A | 19203330 |
| Zld294_GLC_5_1 | Zld294 | 3 | Zld294_FRT19A/Zld294_FRT19A | 21287685 |
| Zld294_GLC_6_1 | Zld294 | 3 | Zld294_FRT19A/Zld294_FRT19A | 19614810 |
| Zld294_GLC_7_1 | Zld294 | 3 | Zld294_FRT19A/Zld294_FRT19A | 21877546 |
| Zld294_GLC_8_1 | Zld294 | 3 | Zld294_FRT19A/Zld294_FRT19A | 20532415 |

**Table S2. Summary of CUT&RUN and ATAC-seq experiments and associated file names submitted to GEO**

| Filename | Condition | Antibody | Assay | Date | Total Reads | % Aligned w/Bowtie2 | Aligned Reads | RiP |
|---|---|---|---|---|---|---|---|---|
| ATAC_H2B | WT | NA | ATAC | 230719 | 28344155 | 87.88 | 24908844 | 0.29 |
| ATAC_Znf5_1 | ZF5 | NA | ATAC | 230719 | 29298587 | 84.07 | 24631323 | 0.35 |
| ATAC_Znf5_2 | ZF5 | NA | ATAC | 230719 | 30503525 | 81.33 | 24808517 | 0.31 |
| ATAC_Zld294_1 | Zld294 | NA | ATAC | 240718 | 10205428 | 89 | 9082831 | 0.08 |
| ATAC_Zld294_2 | Zld294 | NA | ATAC | 240718 | 13155206 | 83.44 | 10976704 | 0.06 |
| ATAC_Zld294_3 | Zld294 | NA | ATAC | 240718 | 14384544 | 84.85 | 12205286 | 0.07 |
| ATAC_Zld294_4 | Zld294 | NA | ATAC | 240718 | 9220031 | 84.74 | 7813055 | 0.09 |
| ATAC_Zld294_5 | Zld294 | NA | ATAC | 240718 | 10205262 | 92.73 | 9463340 | 0.05 |
| ATAC_Zld294_6 | Zld294 | NA | ATAC | 240718 | 15458598 | 90.44 | 13980757 | 0.07 |
| sfGFPzldwt_Gaf1 | WT | Gaf | C&R | 240323 | 7955238 | 16.95 | 1348413 | 0.28 |
| sfGFPzldwt_Gaf2 | WT | Gaf | C&R | 240323 | 11662950 | 14.26 | 1663137 | 0.29 |

| | | | | | | | | |
|---|---|---|---|---|---|---|---|---|
| sfGFPzldwt_Gaf3 | WT | Gaf | C&R | 240704 | 37695502 | 15.25 | 5748565 | 0.33 |
| sfGFPzldwt_Gaf4 | WT | Gaf | C&R | 240704 | 21234142 | 16.58 | 3520621 | 0.35 |
| sfGFPzldwt_Gaf5 | WT | Gaf | C&R | 240704 | 22932052 | 15.52 | 3559055 | 0.14 |
| sfGFPzldwt_Gaf6 | WT | Gaf | C&R | 240704 | 25896729 | 15.52 | 4019173 | 0.15 |
| GLC_mNGzld_Gaf_1 | ZF5 | Gaf | C&R | 231207 | 15290910 | 81.98 | 12535489 | 0.23 |
| GLC_mNGzld_Gaf_2 | ZF5 | Gaf | C&R | 240704 | 15293781 | 69 | 10552709 | 0.15 |
| GLC_mNGzld_Gaf_3 | ZF5 | Gaf | C&R | 240704 | 16422981 | 30.18 | 4956456 | 0.12 |
| GLC_mNGzld_Gaf_4 | ZF5 | Gaf | C&R | 240704 | 23625405 | 28.97 | 6844280 | 0.10 |
| GLC_IgG_1 | ZF5 | IgG | C&R | 230620 | 14094365 | 90.45 | 12748354 | 0.04 |
| GLC_IgG_2 | ZF5 | IgG | C&R | 231207 | 17240038 | 55.5 | 9568222 | 0.04 |
| sfGFPzldwt_zld3 | WT | Zld | C&R | 231207 | 18168704 | 86.64 | 15741366 | 0.07 |
| sfGFPzldwt_zld4 | WT | Zld | C&R | 231207 | 22504068 | 94.11 | 21178579 | 0.14 |
| sfGFPzldwt_zld5 | WT | Zld | C&R | 231207 | 22142523 | 95.7 | 21190395 | 0.15 |
| mNGzld_GLC_1 | ZF5 | Zld | C&R | 230620 | 192874 | 86.89 | 167588 | 0.18 |

| | | | | | | | | |
|---|---|---|---|---|---|---|---|---|
| | | | | | 44 | | 61 | |
| mNGzld_GLC_2 | ZF5 | Zld | C&R | 230620 | 24786487 | 91.14 | 22590405 | 0.08 |
| mNGzld_GLC_3 | ZF5 | Zld | C&R | 230719 | 18340650 | 93.03 | 17062307 | 0.08 |
| mNGzld_GLC_4 | ZF5 | Zld | C&R | 231207 | 19221901 | 73.22 | 14074276 | 0.08 |
| mNGzld_GLC_5 | ZF5 | Zld | C&R | 231207 | 13504968 | 49.79 | 6724124 | 0.19 |
| mNGzld_GLC_6 | ZF5 | Zld | C&R | 231207 | 12479679 | 47.82 | 5967783 | 0.21 |

**Table S3.** Summary statistics of differentially expressed genes (download .csv from supplementary materials)

**Table S4.** Summary statistics for single-molecule data from Fig. 3 (download .csv from supplementary materials)

**Table S5.** Summary statistics of hub properties from Fig. 1 and 4 (download .csv from supplementary materials)

**Table S6.** Summary statistics of hub lifetimes from Fig. 4 (download .csv from supplementary materials)

**Table S7.** Summary statistics of hub-target gene interactions from Fig. 5 (download .csv from supplementary materials)

**Movie Captions**

**Movie 1. Volumetric Imaging of all Zelda mutants in nc13.** 4D volumetric imaging data showing side-by-side comparison of mNeonGreen-ZLD, mNeonGreen-ZF5/+, mNeonGreen-ZF5, mNeonGreen-ΔIDR/+, and mNeonGreen-ZF1-6/+ during interphase of nc13. Scale bar = 5 μm. (Related to Fig. 1).

**Movie 2. Single-Molecule Imaging for determining residence times.** Single molecule data acquired at an exposure time of 500 milliseconds for mEos3.2-His2B, mEos3.2-ZLD, and mEos3.2-ZF5. (Related to Fig. 3).

**Movie 3. Single-Molecule Imaging for quantifying diffusion kinetics of mEos3.2-ZLD.** Single molecule data acquired at an exposure time of 10 milliseconds for mEos3.2-ZLD (Related to Fig. 3).

**Movie 4. Single-Molecule Imaging for quantifying diffusion kinetics of mEos3.2-ZF5.** Single molecule data acquired at an exposure time of 10 milliseconds for meos3.2-ZF5. (Related to Fig. 3).

**Movie 5. Volumetric Imaging of ZLD and ZF5 in nc14.** 4D volumetric imaging data showing side-by-side comparison of mNeonGreen-ZLD and mNeonGreen-ZF5 during interphase of nc14 pre-cellularization. (Related to Fig. 4).

**Movie 6. Volumetric Imaging of ZLD.** 4D volumetric imaging data showing mNeonGreen-ZLD through nc12-14. (Related to Fig. 4).

**Movie 7. Volumetric Imaging of ZF5.** 4D volumetric imaging data showing mNeonGreen-ZF5 through nc12-14. (Related to Fig. 4).

**Movie 8. Volumetric Imaging of sfGFP-ZLD.** 4D volumetric imaging data showing sfGFP-ZF5 in nc14. (Related to Fig. 4).

**Movie 9. Hub Segmentation.** Example of 4D hub segmentation for mNeonGreen-ZLD. Zelda-mNeonGreen is shown in gray, and hub masks are shown in fire LUT (purple/red) (Related to Fig. 1 and 4).

**Movie 10. Volumetric Imaging of ZF5/+.** 4D volumetric imaging data showing mNeonGreen-ZF5/+ through nc12-14. (Related to Fig. 4).

**Movie 11. Volumetric Imaging of ΔIDR/+.** 4D volumetric imaging data showing mNeonGreen-ΔIDR/+ through nc12-14. (Related to Fig. 4).

**Movie 12. Volumetric Imaging of ZF1-6/+.** 4D volumetric imaging data showing mNeonGreen-ZF1-6/+ through nc12-14. (Related to Fig. 4).

**Movie 13. Simultaneous volumetric Imaging of ZLD in the context of nascent transcription of *Antp*.** Volumetric imaging data of mNeonGreen-ZLD (gray) and MCP-mCherry (magenta) localized to

*Antp-MS2* in late nc14. Cytoplasmic background of MCP-mCherry is due to the protein lacking a NLS to maximize signal to noise within nuclei. (Related to Fig. 5).

**Movie 14. Simultaneous volumetric Imaging of ZF5 in the context of nascent transcription of *Antp* in nc12.** Volumetric imaging data of mNeonGreen-ZF5 (gray) and MCP-mCherry (magenta) localized to *Antp-MS2* in nc12. (Related to Fig. 5).

**Movie 15. Simultaneous volumetric Imaging of ZF5 in the context of nascent transcription of *Antp* in nc13.** Volumetric imaging data of mNeonGreen-ZF5 (gray) and MCP-mCherry (magenta) localized to *Antp-MS2* in nc13. (Related to Fig. 5).

**Movie 16. Simultaneous volumetric Imaging of ZF5 in the context of nascent transcription of *Antp* in nc14.** Volumetric imaging data of mNeonGreen-ZF5 (gray) and MCP-mCherry (magenta) localized to *Antp-MS2* in nc14. (Related to Fig. 5).
